# Structurally-informed human interactome reveals proteome-wide perturbations by disease mutations

**DOI:** 10.1101/2023.04.24.538110

**Authors:** Dapeng Xiong, Yunguang Qiu, Junfei Zhao, Yadi Zhou, Dongjin Lee, Shobhita Gupta, Mateo Torres, Weiqiang Lu, Siqi Liang, Jin Joo Kang, Charis Eng, Joseph Loscalzo, Feixiong Cheng, Haiyuan Yu

**Affiliations:** Department of Computational Biology, Cornell University, Ithaca, NY 14853, USA; Weill Institute for Cell and Molecular Biology, Cornell University, Ithaca, NY 14853, USA; Center for Innovative Proteomics, Cornell University, Ithaca, NY 14853, USA; Genomic Medicine Institute, Lerner Research Institute, Cleveland Clinic, Cleveland, OH 44195, USA; Department of Systems Biology, Herbert Irving Comprehensive Center, Columbia University, New York, NY 10032, USA; Biophysics Program, Cornell University, Ithaca, NY 14853, USA; Shanghai Key Laboratory of Regulatory Biology, Institute of Biomedical Sciences and School of Life Sciences, East China Normal University, Shanghai 200241, China; Department of Molecular Medicine, Cleveland Clinic Lerner College of Medicine, Case Western Reserve University, Cleveland, OH 44195, USA; Case Comprehensive Cancer Center, Case Western Reserve University School of Medicine, Cleveland, OH 44106, USA; Channing Division of Network Medicine, Division of Cardiovascular Medicine, Department of Medicine, Brigham and Women’s Hospital, Harvard Medical School, Boston, MA 02115, USA

**Author notes:** Corresponding authors. (F.C.), (H.Y.). These authors contributed equally (Dapeng Xiong, Yunguang Qiu, Junfei Zhao, Yadi Zhou, Dongjin Lee).

## Abstract

Human genome sequencing studies have identified numerous loci associated with complex diseases. However, translating human genetic and genomic findings to disease pathobiology and therapeutic discovery remains a major challenge at multiscale interactome network levels. Here, we present a deep-learning-based ensemble framework, termed PIONEER (**P**rotein-protein **I**nteracti**O**n i**N**t**E**rfac**E** p**R**ediction), that accurately predicts protein binding partner-specific interfaces for all known protein interactions in humans and seven other common model organisms, generating comprehensive structurally-informed protein interactomes. We demonstrate that PIONEER outperforms existing state-of-the-art methods. We further systematically validated PIONEER predictions experimentally through generating 2,395 mutations and testing their impact on 6,754 mutation-interaction pairs, confirming the high quality and validity of PIONEER predictions. We show that disease-associated mutations are enriched in PIONEER-predicted protein-protein interfaces after mapping mutations from ∼60,000 germline exomes and ∼36,000 somatic genomes. We identify 586 significant protein-protein interactions (PPIs) enriched with PIONEER-predicted interface somatic mutations (termed oncoPPIs) from pan-cancer analysis of ∼11,000 tumor whole-exomes across 33 cancer types. We show that PIONEER-predicted oncoPPIs are significantly associated with patient survival and drug responses from both cancer cell lines and patient-derived xenograft mouse models. We identify a landscape of PPI-perturbing tumor alleles upon ubiquitination by E3 ligases, and we experimentally validate the tumorigenic KEAP1-NRF2 interface mutation p.Thr80Lys in non-small cell lung cancer. We show that PIONEER-predicted PPI-perturbing alleles alter protein abundance and correlates with drug responses and patient survival in colon and uterine cancers as demonstrated by proteogenomic data from the National Cancer Institute’s Clinical Proteomic Tumor Analysis Consortium. PIONEER, implemented as both a web server platform and a software package, identifies functional consequences of disease-associated alleles and offers a deep learning tool for precision medicine at multiscale interactome network levels.

## Introduction

Precision medicine has sparked major initiatives focusing on whole-genome/exome sequencing (WGS/WES) and developing genome science tools for statistical analyses – all aspiring to identify actionable variants in patients^1–3^. At the center of the massive amount of collected DNA/RNA sequencing data is their functional interpretation, which largely rests on conventional statistical analyses and trait/phenotype observations^2^. Statistics is vital, since it guides the identification of disease-associated variants; however, traditional WGS/WES studies are commonly underpowered for disease risk-variant/gene and drug target discoveries as very large sample sizes are generally required. Furthermore, the statistics do not directly elucidate the functional consequence of the variants, i.e., changes in protein conformation or in molecular functions in cells; thus, traditional statistical approaches may not be sufficient to define accurately functional variants. Thus, translation of genetic and genomic findings to precision medicine is fraught with challenges using traditional statistical approaches.

Optimal information requires knowledge of the whole protein-protein interaction (PPI) network, or interactome, within which the mutant protein operates. Given that on average each protein interacts directly with 10-15 other proteins^4, 5^, the functional consequence of any mutation is not easily (if at all) predicted out of the interactome context. Previous studies^3, 6–10^ have demonstrated that most disease mutations disrupt specific PPIs rather than affecting all interactions in which the mutant protein is engaged. Making accurate characterizations of such disruptions is essential for understanding the etiology of most disease mutations. Therefore, it is fundamentally important for precision medicine to determine structural details, particularly the locations of interaction interfaces of all protein interactions at whole proteome scale. A clear limitation of this goal, however, is that only ∼9% of protein interactions have structural models determined by experimental or traditional homology modeling approaches (Figs. 1a and 2a). Predicting co-complex structures of PPIs is in the process of explosive growth resulting from the advent of AlphaFold-based deep learning methods as embodied in AlphaFold-Multimer^11^ or AF2Complex^12^, but these methods are all time-consuming, and does not scale to solve whole interactomes with hundreds of thousands of PPIs.

**Fig. 1.**
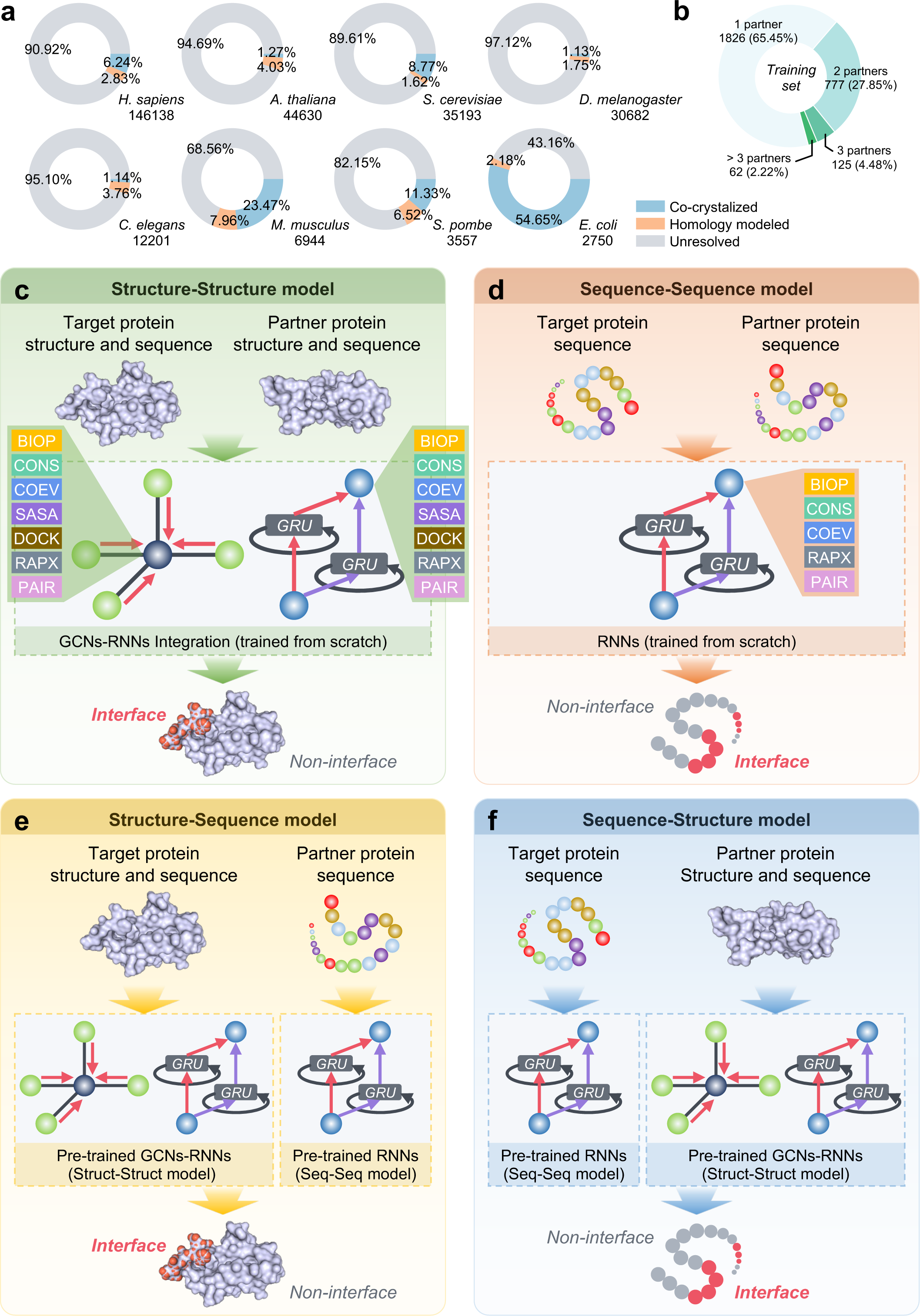
Overview of the PIONEER framework. **a,** The current size of protein-protein interactions from the eight common model organisms with the coverage of experimentally determined co-crystal structures, homology models, and the unresolved interactions. **b,** The partner-specific interactions are prioritized in our training dataset for solving partner-specific interface prediction. **c**-**f**, PIONEER architecture consists of an ensemble of four deep learning models that ensures every residue in the interactomes can be predicted with the maximal amount of available information, and uses a comprehensive set of biophysical, evolutionary, structural, and sequence features for in-depth feature characterization. The **c** and **d** models represent interactions in which both proteins and neither protein has structural information available, respectively. The GCNs and RNNs are used for structure and sequence information embeddings, respectively. The **e** and **f** models represent interactions in which only one protein has structure information available. The transfer learning was used in **e** and **f**, specifically, the pre-trained GCNs-RNNs in **a** and RNNs in **b** are deployed in **c** and **d**.

**Fig. 2.**
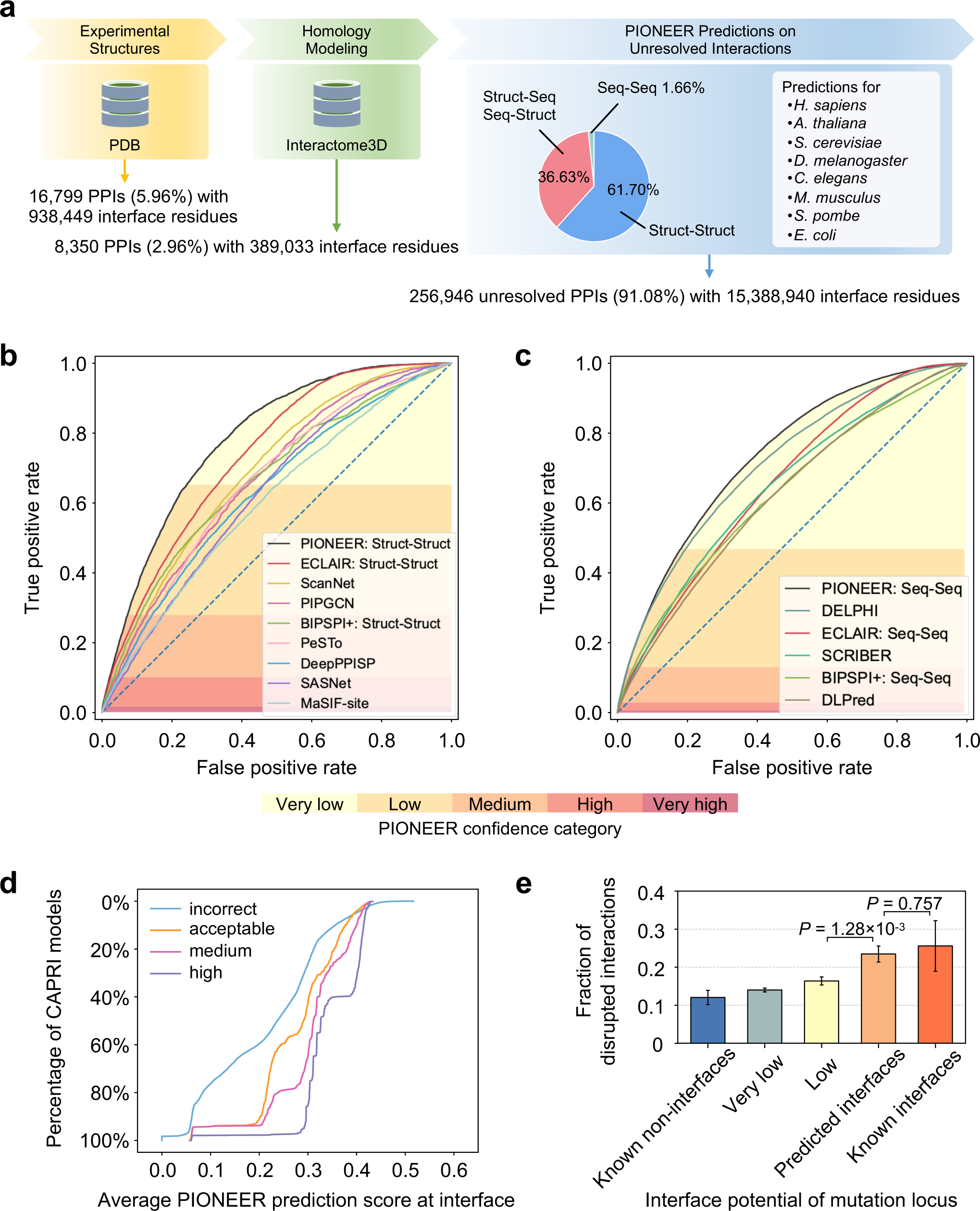
PIONEER provides high-quality interfaces for the whole proteome. **a,** Workflow for compiling interactome PIONEER. The interfaces calculated from experimentally determined co-crystal structures or homology models are primarily used, the remaining unresolved interactions are predicted by PIONEER. **b,** Comparison of receiver operating characteristic (ROC) curves of PIONEER Structure-Structure model with other state-of-the-art structure-based methods. **c,** Comparison of ROC curves of PIONEER Sequence-Sequence model with other state-of-the-art sequence-based methods. **d,** Percentage of CAPRI decoys having a given average PIONEER prediction score at interfaces. **e,** Fraction of interactions disrupted by random population variants in PIONEER-predicted and known interfaces.

In this study, we presented a deep-learning-based ensemble learning pipeline, PIONEER (**P**rotein-protein **I**nteracti**O**n i**N**t**E**rfac**E** p**R**ediction) to generate the next-generation partner-specific interaction interface predictions for all experimentally-determined human PPIs in the literature, providing key structural information for these interactions. Taking as input a pair of interacting proteins, PIONEER effectively identifies the residues that conform the interface of the interaction. By leveraging the available atomic-resolution co-crystal structures along with homology models, we established a comprehensive multiscale structurally-informed human interactome, which consists of 282,095 interactions from humans and seven other commonly studied organisms, including all 146,138 experimentally-determined PPIs for 16,232 human proteins (Figs. 1a and 2a). Through this resource, we investigated the network effects of disease-associated mutations at amino acid resolution within the macromolecular interactome of PPI interfaces. We further explored the widespread perturbations of PPIs in human diseases and their significant impact on disease prognosis and drug responses. This newly constructed structurally-informed interactome database is then combined with disease-associated mutations and functional annotations to create an interactive, dynamic web server (https://pioneer.yulab.org) for genome-wide functional genomics studies. It also allows the users to submit a list of interactions and retrieve the predicted interfaces of the respective interactions using the PIONEER framework. Furthermore, we have converted the PIONEER framework into a software package that is available to the wider community to further help accelerate biological research.

## Results

### A hybrid deep-learning architecture for protein-protein interface prediction

To date, taking into account the structural models of protein interactions—both experimentally-determined and homologically-predicted--we find that an overwhelming majority of interactions (∼91%) still do not have reliable structural information (Fig. 1a). With this key limitation in mind, we built the PIONEER pipeline to generate accurate partner-specific protein-protein interface predictions for protein interactions that currently lack structural information. We carefully constructed our labeled dataset for training, validating, and testing of our classifiers (Supplementary Data 1). When building PIONEER, we especially prioritize instances where the same protein interacts with multiple interaction partners using distinct interfaces in our labeled dataset in order to create a model that better predicts partner-specific interfaces (Fig. 1b). We also require that there are no homologous interactions between any of the two datasets to guarantee the robustness and generalization of our models, and a fair performance evaluation.

We used a comprehensive set of single-protein and interaction-partner-specific features for interface prediction (Fig. 1c-f), and both groups of features combine biophysical, evolutionary, structural, and sequence information for an in-depth feature characterization. More specifically, the single-protein features consist of diverse biophysical features, evolutionary sequence conservation, and protein structure properties. Of these features, one of protein structure properties, solvent accessible surface area (SASA), is widely reported to be highly informative^6, 13^ since all interface residues, by definition, are on the surface of proteins; in addition, solvent can either facilitate or impede binding interactions depending upon the (net) relative hydrophobicity or hydrophilicity of the binding interaction residues. We also incorporate evolutionary sequence conservation because key protein functions often depend upon interactions with other proteins, and, as such, interface residues are more likely to be conserved^14^.

However, although these single-protein features capture the characteristics of all possible interface residues, they cannot distinguish interface residues for a protein interacting with different partner proteins through which a protein can perform different biological functions. Previously, we illustrated that it is important to encompass partner-specific features for the prediction of partner-specific interfaces^6^. Here, our interaction-partner-specific features include co-evolution of amino acid sequences, protein-protein docking, and pair potential. Co-evolutionary features capture the dependent patterns of conservation in two interacting proteins, since interface residues that are critical to maintaining the interaction often co-evolve to maintain binding complementarity^15^. Through modeling of bound conformations of protein interactions using protein-protein docking^16^, we incorporate the docking results to extract a summary of preferred orientations for individual structures of interacting proteins. In addition, we also create another feature—pair potential—to characterize the interface propensity for the target protein with respect to a specific partner. Moreover, we incorporate AlphaFold2-predicted single protein structures into our PIONEER models to significantly increase the coverage of structure-based features for proteins currently lacking experimentally-determined structures.

In order to address the non-random missing feature problem, which cannot be adequately resolved by commonly used imputation methods^6^, PIONEER’s framework is structured as an ensemble of four deep learning architectures, including Structure-Structure, Structure-Sequence, Sequence-Structure, and Sequence-Sequence models (Fig. 1c-f and Supplementary Figs. 1 and 2). The Structure-Structure model is used for interactions in which both proteins have structural information while the Sequence-Sequence model is used for proteins without solved structural information. Otherwise, the Structure-Sequence or Sequence-Structure model is used, depending on which protein in the interaction has structural information. This comprehensive framework ensures that we use the maximum amount of information available for each interaction to yield the best possible interface predictions while avoiding potential ascertainment biases that can lead to overfitting.

More specifically, for a protein with available structures, the PIONEER deep learning model uses a hybrid architecture to integrate both structural information embedded through Graph Convolutional Networks (GCNs) with Auto-Regressive Moving Average (ARMA) filters^17^ and sequence information embedded through bidirectional Recurrent Neural Networks (RNNs) with Gated Recurrent Units (GRUs)^18^. For proteins without high-quality structure models, only sequence information is embedded via RNNs with GRUs. Using transfer learning^19^, the pre-trained GCNs-RNNs in Structure-Structure model and RNNs in Sequence-Sequence model are deployed in Structure-Sequence model and Sequence-Structure model for the processing of proteins with and without structural information, respectively. Furthermore, for each residue in a target protein, our unique architecture integrates the residue embeddings, overall protein embeddings, and overall partner protein embeddings to utilize both local and global (‘glocal’) information of both proteins to make the most accurate interface predictions.

### Benchmark evaluation of PIONEER

We then built and evaluated PIONEER using the new labeled dataset and new model designs. The results show that PIONEER outperforms all other available methods for predicting interactions of both proteins with and without structural information (Fig. 2b,c and Supplementary Figs. 3 and 4 and Supplementary Tables 1-5). It is worth noting that our Sequence-Sequence model, which solely relies on sequence information, has better prediction performance than all recent state-of-the-art structure-based methods that we evaluated, such as PeSTo^20^, ScanNet^21^, BIPSPI+^22^, MaSIF-site^23^, DeepPPISP^24^, SASNet^25^, and PIPGCN^26^. Most of these methods already use cutting-edge deep learning models, which illustrates the power of utilizing a comprehensive set of single-protein and partner-specific features; it also confirms the validity of our design choice to include RNNs with GRUs in a hybrid architecture, even for proteins with known structures (Fig. 2b and Supplementary Tables 2 and 3). Interestingly, we also found that even our previous ECLAIR with structural information is still significantly better than the above structure-based methods, and achieves the second best performance (Fig. 2c and Supplementary Table 2).

We next evaluated the effectiveness of our new model designs on the benchmark testing dataset by assessing the overall performance of PIONEER and ECLAIR. We found that PIONEER models with ECLAIR features substantially outperform our previous ECLAIR models (Supplementary Fig. 5a), confirming that our unique hybrid-architecture deep learning models, indeed, capture more information in the features than the previous random forest-based models. Moreover, incorporating new features to PIONEER models further significantly improves the prediction performance (Supplementary Fig. 5a), indicating the outstanding representation ability of our new features for the prediction of protein interfaces. Both improvements distinctly demonstrate that our new deep learning architectures and new features make significant contributions to PIONEER’s strong ability to provide accurate PPI interface predictions. We also found that the inclusion of AlphaFold2-predicted single protein structures for the proteins without experimentally-determined structures improved the performance of PIONEER interface predictions (Supplementary Fig. 5b).

We further applied PIONEER on a widely used decoy set, score_set^27^, to test CAPRI (Critical Assessment of Prediction of Interactions) models. This dataset contains docking models predicted by 47 different groups for proteins from bacteria, yeast, vertebrates, and artificial design. After removing the targets which are duplicated and without corresponding UniProt sequences, we considered 11 targets which, overall, have 15,003 corresponding decoys, including 12,986 incorrect, 732 acceptable, 799 medium and 486 high quality docking predictions based on CAPRI-defined criteria, respectively. Figure 2d plots the percentage of all models that have a given average PIONEER interface probability score (x-axis) in each category across targets. This figure shows a clear distinction between any two different category decoys, demonstrating that PIONEER interface residue predictions also provide a clear signal as to model quality.

### Proteome-wide protein-protein interface prediction by PIONEER

Next, we compiled a comprehensive set of experimentally validated binary PPIs for humans and seven model organisms by integrating information from 7 commonly-used databases^28^, including BioGRID^29^, DIP^30^, IntAct^31^, MINT^32^, iRefWeb^33^, HPRD^34^, and MIPS^35^. Here we focus on binary interactions because if two proteins do not bind to each other directly, the concept of interface does not apply. We then used the fully-optimized PIONEER pipeline to predict interfaces for all collected 256,946 binary interactions without any experimental structures or homologous models (see Methods), including 132,875 human interactions (Fig. 2a). Because we make partner-specific interface predictions for every residue of every protein, and there are on average >10 interactions per protein, we made probabilistic interaction predictions for >275 million residue-interaction pairs. By combining our PIONEER interface predictions with 25,149 interactions (∼9%) with experimental or homology models, we generate a comprehensive multiscale structural human interactome, in which all interactions have partner-specific interface information at the residue level, together with atomic-resolution three-dimensional (3D) models whenever possible.

To comprehensively evaluate the quality of our predicted interfaces and their biological implications, we first carried out large-scale mutagenesis experiments to measure the fraction of disrupted interactions by mutations in our predicted interfaces at varying confidence levels, in comparison to that of known interface and non-interface residues from co-crystal structures in the PDB^36^. Using our Clone-seq pipeline^37^, we generated 2,395 mutations on 1,141 proteins and examined their impact on 6,754 mutation-interaction pairs through a high-throughput yeast-two-hybrid (Y2H) assay, an unprecedented large-scale experimental validation. We observed that mutations at PIONEER-predicted interfaces disrupt protein-protein interactions at a very similar rate to the mutations at known structurally characterized interfaces, and both of their disruption rates are significantly higher than that of known non-interfaces (Fig. 2e). Therefore, our large-scale experiments confirm the high quality of our interface predictions and the validity of our overall PIONEER pipeline. Furthermore, because interaction disruption is key to understanding the molecular mechanisms of disease mutations^9, 37^, our experimental results indicate that our PIONEER-predicted interfaces will be instrumental in prioritizing disease-associated variants and generating concrete mechanistic hypotheses.

### PIONEER-generated structurally-informed human interactome enriched with disease mutations

Since disruption of specific interactions is essential for the pathogenicity of many disease mutations, and since previous studies have shown that disease mutations are significantly enriched at protein-protein interfaces^7, 8, 38^, we next measured the enrichment of known disease-associated mutations from the Human Gene Mutation Database (HGMD)^39^ at the predicted interfaces, and compared it to known interfaces from co-crystal structures. We found that the residues predicted by PIONEER with a high interface confidence show a very similar rate of disease mutation enrichment to those of known interfaces (Fig. 3a). Furthermore, we observed that 251,368 (∼98%) out of all 256,946 binary interactions have at least one or more predicted interface residues that fall into high or very high confidence categories (Supplementary Fig. 6), indicating that PIONEER provides meaningful structural information for the vast majority of protein interactions. In fact, each bin with a higher confidence of predicted interfaces is more likely to contain disease-associated mutations than the previous bin, which demonstrates the strong correlation between PIONEER prediction scores and the true function of each residue (Fig. 3a). We further analyzed the distribution of population genetic variants and found that their enrichment in PIONEER-predicted interfaces and non-interfaces also matches well with that of known interfaces and non-interfaces, respectively (Fig. 3b). The results also show that there is a depletion of common variants (i.e., not deleterious) in both known and predicted interfaces, which strongly suggests that there exists a strong negative selection at these interface residues, indicating that PIONEER is able to predict functionally important interface variants effectively. We also found that, compared to variants identified in individuals from the 1000 Genomes project (1KGP)^40^ and the Exome Aggregation Consortium (ExAC)^41^, disease-associated mutations from HGMD are more significantly enriched at PPI interfaces of the respective proteins^8^ (Fig. 3c). Moreover, as predicted by CADD^42^ and FoldX^43^, the population variants in PIONEER-predicted interfaces are more likely to adversely affect protein functions than those in PIONEER-predicted non-interfaces (Supplementary Fig. 7), which confirms that deleterious variants preferentially occur at the protein-protein interfaces^7, 9^.

**Fig. 3.**
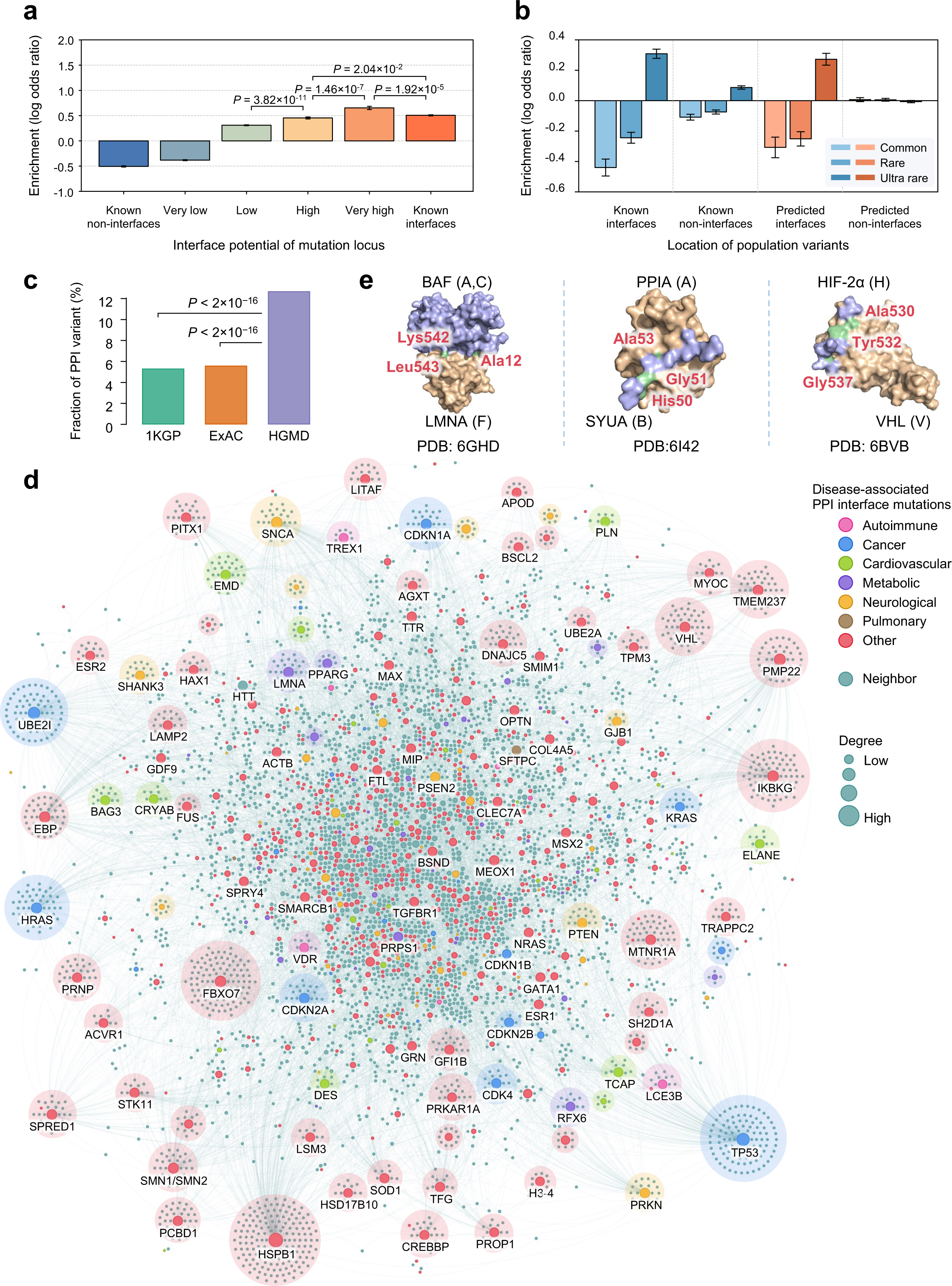
PIONEER-predicted PPI alleles are enriched in disease-associated mutations. **a,** Enrichment of disease mutations in PIONEER-predicted and known interfaces. Significance was determined by two-tailed Z-test. **b,** Enrichment of population variants in PIONEER-predicted and known interfaces. Predicted deleteriousness of population variants in PIONEER-predicted and known interfaces using PolyPhen-2. **c,** Distribution of mutation burden at protein–protein interfaces for disease-associated germline mutations from HGMD in comparison with mutations from the 1KGP and ExAC. Significance was determined by two-proportion Z-test. **d,** PPI network with disease-associated interface mutations. Disease associations of the interface mutations were extracted from the HGMD database. Using the PIONEER-predicted high-confidence interface information, PPIs that have at least one disease-associated interface mutation from either one of the two interacting proteins were included in the network. Node colors were determined by the disease categories of their disease-associated interface mutations. Interacting proteins with no known disease-associated interface mutations were colored as “Neighbor”. The final network contains 10,753 PPIs among 5,684 proteins. The figure shows the largest connected component of the network that has 10,706 edges and 5,605 nodes. **e,** Selected structural complex pairs showing germline mutations in the PPI interface. Three disease associated PPIs with mutations are shown: LMNA-BAF (PDB: 6GHD), PPIA-SYUA (PDB: 6I42), and VHL-HIF-2α (PDB: 6BVB). The surface of proteins in complex are shown in blue and wheat. Mutations are shown in green.

To further evaluate whether the disease-associated mutations were enriched in the PIONEER-predicted PPI interfaces (Fig. 3c), we next categorized the disease-associated germline mutations from HGMD into six major disease groups, including autoimmune, cancer, cardiovascular, metabolic, neurological, and pulmonary as described in a recent study^44^, as well as an additional “other” category. We identified a total of 10,753 PPIs among 5,684 proteins that had at least one disease-associated interface germline mutation (Fig. 3d and Supplementary Table 6), among which 9,795 (∼91%) PPIs have such interface mutations on one protein (the other protein colored as “neighbor”), and 958 (∼9%) on both proteins of the binding pair. Overall, this network analysis shows that PIONEER-predicted PPI interfaces are altered by broad disease-associated mutations across multiple disease categories. To highlight the power of PIONEER-predicted interfaces, we examined three PPI interfaces with germline alleles. Germline mutation p.Lys542Gln of LMNA (Lamin A/C) buried in the interface of LMNA and BAF (Fig. 3e) was associated with progeroid disease^45^. One loss-of-function PPIA mutation p.Ala53Glu in the interface of PPIA-SYUA (Fig. 3e) was identified in patients with early-onset Parkinson disease^46^. A germline mutation p.Gly537Arg of HIF-2α associated with polycythemia vera^47^ is located in the PIONEER-predicted VHL-HIF-2α interface (Fig. 3e) and disrupts VHL binding via impairing ubiquitination and proteasomal degradation of HIF-2α^48, 49^. Taken together, PIONEER-predicted protein-protein interface mutations convey crucial structural information in delineating the functional consequences for disease mechanisms at the atomic and allele levels.

### PIONEER-predicted oncoPPIs across 33 cancer types

We next investigated the somatic mutations from cancer patients in the context of PPI interfaces inferred by PIONEER. In total, we collected ∼1.7 million missense somatic mutations from the analysis of ∼11,000 tumors across 33 cancer types from The Cancer Genome Atlas (TCGA)^50^. We found significant enrichment of somatic missense mutations in the PIONEER-predicted PPI interfaces compared to non-interface regions (Fig. 4a and Supplementary Data 2). Specifically, this significant enrichment was observed in 31 out of the total 33 cancer types regardless of the overall mutation burden. In lung squamous cell carcinoma (LUSC), one of the cancer types that have the highest mutation load per exome, we observed 29 variants per 1 million amino acid residues affecting PPI interfaces, while the rate for the non-PPI interface region is 23 (*P* = 1.3×10^−11^). For thyroid cancer (THCA), with the lowest mutation load, the difference is 27 for PPI interfaces vs. 9 for the remainder of the protein sequences (*P* < 10^−16^). To account for the potential bias in this analysis due to data sources, we divided our whole structural human interactome into three categories: those with experimental structures (PDB, 6.2%), with homology models (HM, 2.8%), and predicted by PIONEER (90.9%), and performed the enrichment analysis for each category separately. The results showed that the same enrichment pattern we observed is independent of the data source, suggesting the robustness of PIONEER interface predictions (Supplementary Fig. 8).

**Fig. 4.**
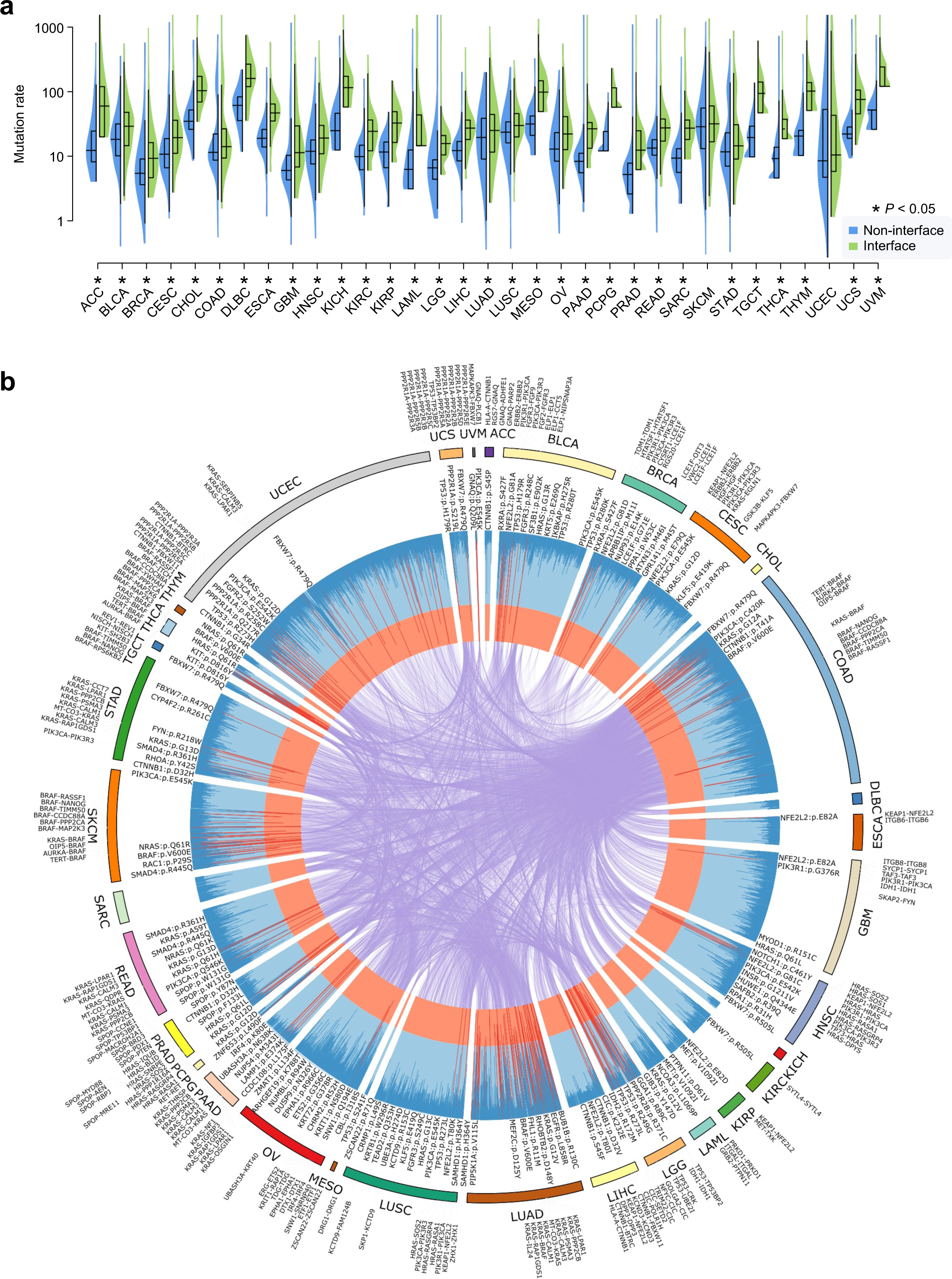
A landscape of oncoPPIs identified by PIONEER across 33 cancer types (∼11,000 cancer genomes). **a,** Distribution of missense somatic mutations in protein–protein interfaces versus non-interfaces across 33 cancer types/subtypes from TCGA. The data are represented as violin plots with underlaid boxplots, where the middle line is the median, the lower and upper edges of the rectangle are the first and third quartiles, and the lower and upper whiskers of the violin plot represent the interquartile range (IQR) × 1.5. Significance was determined by two-tailed Wilcoxon rank-sum test. **b,** Circos plot displaying significant putative oncoPPIs harboring a statistically significant excess number of missense somatic mutations at PPI interfaces across 33 cancer types. Putative oncoPPIs with various significance levels are plotted in the three inner layers. The links (edges, orange) connecting two oncoPPIs indicate two cancer types sharing the same oncoPPIs. Selected significant oncoPPIs and their related mutations are plotted on the outer surface. The length of each line is proportional to –log_10_(*P*).

We next sought to identify the oncoPPIs that were significantly enriched by somatic mutations in the interfaces in both individual cancer types and pan-cancer analyses. Specifically, the average number of PPIs affected by interface mutations is 10,741. Finally, our analysis yielded total 586 statistically significant oncoPPIs across 33 cancer types (Fig. 4b and Supplementary Data 3), including KRAS-BRAF, TP53-EGLN1, and TP53-TP53BP2 across 10 cancer types.

We next turned to analyze the clinical sequencing data from MSK-MET, a pan-cancer cohort of over 25,000 patients spanning 50 different tumor types^51^. Of the 157,979 missense mutations we investigated, 40,526 (∼26%) were identified to affect 15,523 unique PPI interfaces. Focusing on the PPIs that were disturbed in at least 10 samples in a specific cancer type, we performed survival analysis to identify clinically actionable oncoPPIs whose disruption is significantly associated with patient survival. KRAS has been reported to co-mutate with NF1 (neurofibromin protein) in response to GTP hydrolysis^52^. We identified that the mutations of KRAS-NF1 interface residues, such as Asp30 and Glu31 on KRAS (Fig. 5a), are significantly associated with poor survival rate compared to the wild-type (WT) group in pancreatic cancer (*P* = 2.7×10^−18^; Fig. 5b). SPOP plays a multifaceted role in oncogenesis and progression by mediating degradation of PTEN^53^, BRD3^54^, TP53BP1^55^, PDX1^56^, and MACROH2A1^57^. The SPOP MATH domain binds to PTEN via a PIONEER-predicted interface mutation of p.Phe133Val on SPOP^58^, which is significantly associated with survival rate in prostate cancer (*P* = 0.0021; Fig. 5b). Patients with several PIONEER-predicted interface mutations (Thr231, Pro191 and Arg181 on TP53; Fig. 5a) between TP53 (a key tumor suppressor gene) and KDM4D (a histone demethylase) are significantly associated with poor survival in soft tissue sarcoma (*P* = 0.031; Fig. 5b). OncoPPI analysis revealed that PIONEER-predicted interface mutations in ARIH2-TP53, kinase-substrate (e.g., KIT-BLK), kinase-E3 ligase (e.g., MAPKAPK3-FBXW7), and cyclin-E3 ligase (e.g., CCND1-FBXO31), are significantly associated with survival rate in breast cancer (*P* = 1 × 10^−4^), gastrointestinal stromal tumor (*P* = 0.011), non-small cell lung cancer (*P* = 0.012), and endometrial cancer (*P* = 0.024), respectively (Fig. 5b and Supplementary Fig. 9). Accumulated evidence suggested the mutations of CCND1 are associated with multiple cancer types^59^. By analyzing PIONEER-predicted oncoPPIs, we found PIONEER-predicted interface mutations of CCND1 are significantly enriched at the CCND1-CDK4 interfaces in uterine cancer (*P* = 0.012) and low-grade glioma (*P* = 0.048). We identified that CCND1 not only interacts with CDK4 but also TSC2 from PIONEER-predicted interfaces. Specifically, we identified that CCND1 interacts with CDK4 and TSC2 via two unique sets of interfaces (Fig. 5c). Next, we experimentally confirmed this result using co-immunoprecipitation with 293T cells. Fig. 5d shows that a mutation of p.Lys114Arg on CCND1 specifically disrupts the interaction between CCND1 and CDK4, without disrupting its interaction with TSC2. Interestingly, mutation p.Glu162Lys on CCND1 does not disrupt its interaction with CDK4, but does disrupt its interaction with TSC2. Both p.Lys114Arg and p.Glu162Lys on CCND1 are associated with myeloma^60^ and lung cancer^61^, respectively. These results further demonstrate that PIONEER-generated structural human interactome can uncover tumorigenesis with distinctive functions corresponding to distinct interfaces, even for those on the same proteins.

**Fig. 5.**
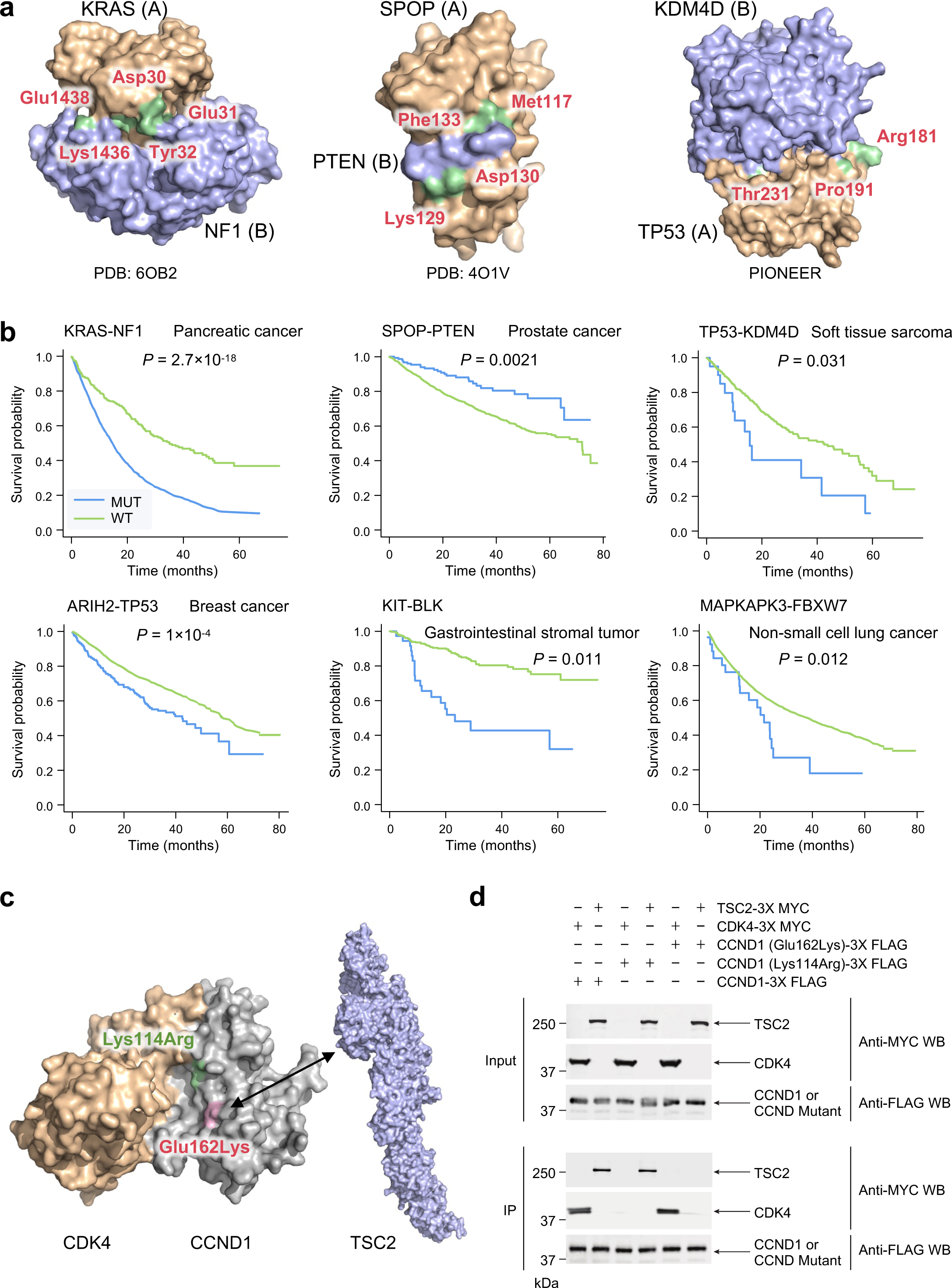
PIONEER-predicted oncoPPIs are associated with patient survival. **a,** Selected structural complex pairs showing somatic mutations in the oncoPPI interface. **b,** Survival analysis of six exemplary PPI perturbing mutations in diverse cancer types. Significance was determined by log-rank test. **c,** Example of PIONEER partner-specific interface prediction. The mutations CCND1 p.Lys114Arg and CCND1 p.Glu162Lys are shown in green and pink, respectively. **d,** Experimental validation of the partner-specific interface prediction in **c** by co-immunoprecipitation using HEK293T cells.

### PIONEER-predicted PPI-perturbing tumor alleles alter ubiquitination by E3 ligases

E3 ubiquitin ligases (E3s) are involved in cellular transformation and tumorigenesis by targeted protein degradation^62, 63^. Identifying how somatic mutations alter PPIs of E3 ligases may offer novel targets for development of targeted protein degradation therapies^64^. We investigated 4,614 PIONEER-predicted oncoPPIs connecting 355 E3 ligases annotated from E3net^65^ and Ubinet2.0^66^ databases. We next focused on the 204 putative oncoPPIs connecting E3 ligases using subject matter expertise based on a combination of factors: i) strong prediction from oncoPPI mutations on E3 ligases or their substrates; ii) significant association with patient survival rates; and iii) significant association with drug responses measured in tumor cell lines or patient-derived tumor xenograft (PDX) mouse models (Supplementary Table 7).

Fig. 6a illustrates the selected examples of the most significant PIONEER-predicted oncoPPIs of E3 ligases (Supplementary Table 8). Among 204 putative oncoPPIs of E3 ligases, FBXW7 (F-box with 7 tandem WD40) has the greatest number of oncoPPIs (47/204; Supplementary Fig. 10 and Supplementary Table 9). FBXL17 (F-box/LRR-repeat protein 17) is a multiple-RING E3 ligase that specifically recognizes and ubiquitinates the BTB proteins^67^. We found that PIONEER-predicted PPI-perturbing mutations of FBXL17-KEAP1, such as p.Ser102Leu on KEAP1, were significantly associated with poor survival in non-small cell lung cancer (*P* = 1.6 × 10^−13^; Fig. 6b). A multiple-RING E3 ligase complex ANAPC1-ANAPC2 (Fig. 6a) is positively regulated by the PTEN/PI3K/AKT pathway and modulates ubiquitin-dependent cell cycle progression^68^. We found that PIONEER-predicted PPI-perturbing mutations on ANAPC1-ANAPC2 is associated with resistance to a PI3K inhibitor BKM120 (*P* = 0.0043; Supplementary Fig. 11a). ITCH, a HECT-type E3 ubiquitin ligase, has been reported to mediate BRAF kinase poly-ubiquitination and promote proliferation in melanoma cells^69^. We found that PIONEER-predicted PPI-perturbing mutations of BRAF-ITCH, such as p.Val600Glu and p.Lys601Glu on BRAF, are significantly associated with sensitivity to dabrafenib (an ATP-competitive inhibitor; *P* = 1.7 × 10^−21^; Supplementary Fig. 11b). STUB1, a U-box-dependent E3 ubiquitin ligase, has been reported to degrade SMAD4, an intracellular signaling mediator of the TGF-β pathway^70^. Multiple PIONEER-predicted PPI-perturbing mutations of STUB1-SMAD4, including p.Gly419Arg (Trp, Val) and p.Leu540Pro (Arg) on SMAD4, are significantly associated with poor survival rate in colorectal cancer (*P* = 0.025; Supplementary Fig. 11c). A single-RING E3 ligase TRIM24 is an oncogenic transcription cofactor overexpressed in breast cancer^71^. We found that PIONEER-predicted PPI-perturbing mutations of TRIM24-H3C1 (Fig. 6a) are significantly associated with resistance to GDC0941 (an EGFR signaling inhibitor; *P* = 0.028; Supplementary Fig. 11d). Treatment with an EGFR inhibitor suppresses TRIM24 expression and H3K23 acetylation and thereby inhibits EGFR-driven tumor growth^72^, supporting the PIONEER-predicted oncoPPI findings.

**Fig. 6.**
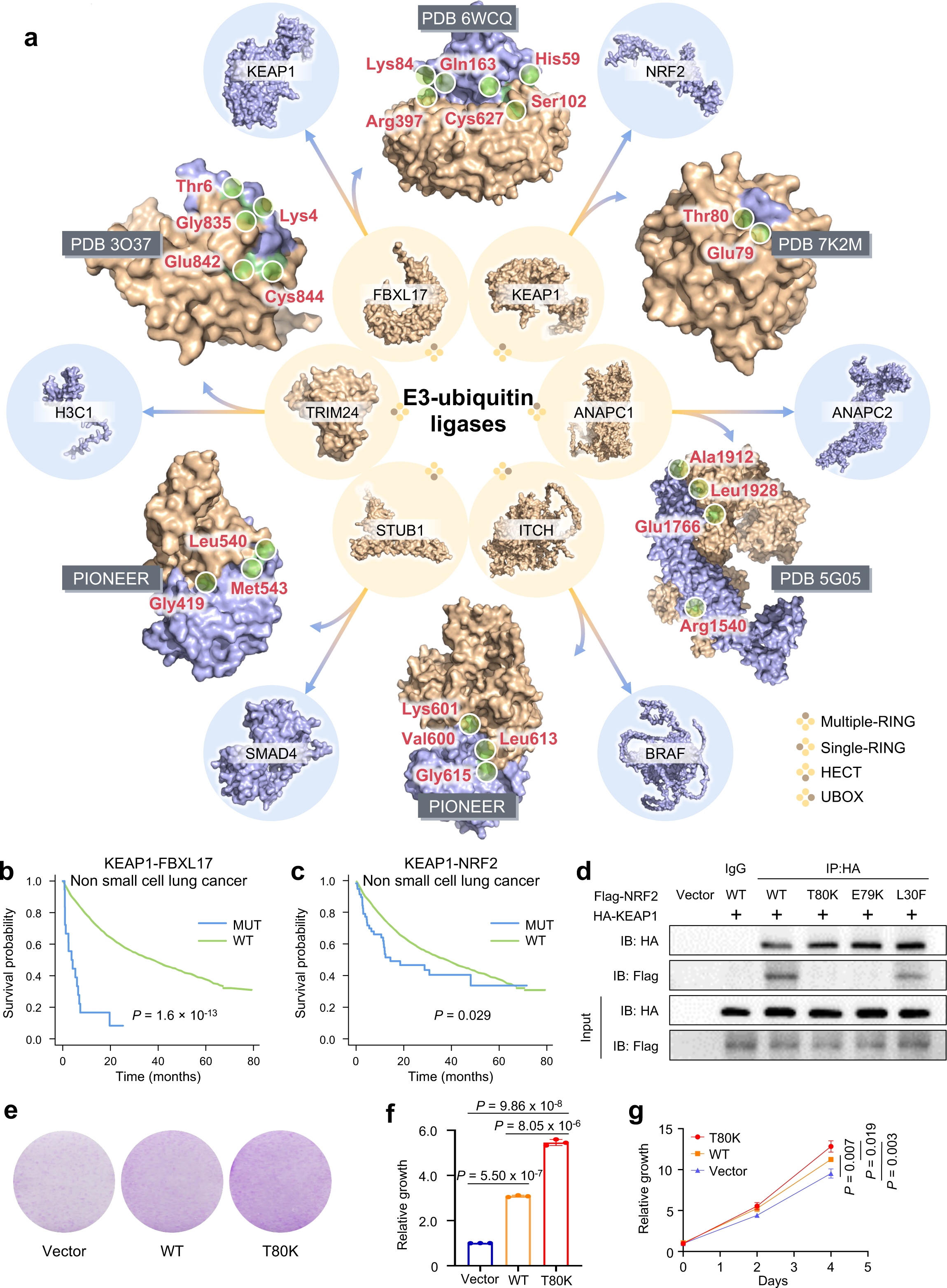
PIONEER-predicted PPI-perturbing tumor alleles in ubiquitination by E3 ligases. **a,** A landscape of six E3 complexes with PPI perturbing mutations. The complex or single protein models from PDB or PIONEER modeling are shown. The protein in wheat denotes the E3 ligase, while the protein in blue denotes the specific substrate of E3 ligase. Interface mutations are denoted in green. **b** and **c,** Interface mutations of KEAP1-FBXL17 (**b**) and KEAP1-NRF2 (**c**) are significantly correlated with survival rate in non-small cell lung cancer. Significance was determined by log-rank test. **d,** Experimental validation of mutation effects of p.Thr80Lys and p.Glu79Lys on the NRF2 ETGE motif, and p.Leu30Phe on the NRF2 DLG motif on the interactions between KEAP1 and WT NRF2 were determined by co-immunoprecipitation with HEK293T cells. **e and f,** Colony formation assay of H1975 cells transfected with NRF2 p.Thr80Lys, NRF2 WT expressing vectors, and empty vectors. **g,** Growth curves of H1975 cells transfected with NRF2 p.Thr80Lys, NRF2 WT expressing vectors, and empty vectors, at day 4.

KEAP1 is an adapter of E3 ligase that senses oxidative stress by mediating degradation of NFE2L2/NRF2, a key transcription factor in multiple cancer types^73^. Patients with non-small cell lung cancer harboring PIONEER-predicted oncoPPI mutations on NRF2 have significantly worse survival than the WT (*P* = 0.029; Fig. 6c). KEAP1 recognizes NRF2 structurally through its conserved ETGE (aa 79-82) and DLG (aa 29-31) motifs^74, 75^. We experimentally confirmed the association of NRF2 mutations and WT KEAP1 by Co-immunoprecipitation. As shown in Fig. 6d, p.Glu79Lys and p.Thr80Lys mutations on the NRF2 ETGE motif (Fig. 6a) reduce the binding of NRF2 to KEAP1, whereas the p.Leu30Phe mutation on the NRF2 DLG motif partially sustains the binding of NRF2 to KEAP1. The mutation of p.Thr80Lys releases NRF2 from association with KEAP1 and protects NRF2 from ubiquitination and subsequent degradation. We next tested whether p.Thr80Lys on NRF2 contributes to the proliferation of non-small cell lung cancer cells. A pro-proliferative effect of p.Thr80Lys was observed in a colony formation assay (Fig. 6e,f). Overexpression of WT and p.Thr80Lys NRF2 promoted the growth of the non-small cell lung cancer H1975 cell line harboring WT KEAP1 (Fig. 6g). In summary, PIONEER-predicted oncoPPI-perturbing tumor alleles that alter ubiquitination by E3 ligases are significantly associated with patient survival, drug responses, and *in vitro* tumor growth.

### Pharmacogenomic landscape of the PIONEER-predicted oncoPPIs

We next turned to inspect correlation between potential oncoPPIs and drug responses using high-throughput drug screening data (Fig. 7a). The datasets we used include the drug pharmacogenomic profiles of >1,000 cancer cell lines and ∼250 FDA-approved or clinical investigational agents from the Genomics of Drug Sensitivity in Cancer (GDSC) database and *in vivo* compound screens using ∼1,000 PDX models to assess patient responses to 62 anticancer agents^76^. For each pair of oncoPPI and compound, the drug response characterization IC_50_ vector was correlated with the mutation status of the oncoPPIs using a linear ANOVA model. Fig. 7b shows the landscape of the correlations between PPIs and 56 clinically investigational or approved anti-cancer drugs. In total, we identified 4,473 interface mutations that have significant correlations with drug sensitivity/resistance. Among the most significant correlations from PDX models, we found that PIONEER-predicted CDK6-BECN1 interface mutations was associated with resistance to treatment using a BYL719 plus encorafenib drug combination, while the mutations in PIONEER-predicted BRAF-MAP2K3 interfaces (e.g., interface mutations p.Val600Glu on BRAF and p.Arg152Gln on MAP2K3, both found in bladder urothelial carcinoma and glioblastoma) conferred significant drug sensitivity to encorafenib plus binimetinib treatment (Fig. 7c). In addition, we found significant drug resistance to trastuzumab and BYL719 among those cases harboring mutations in PIONEER-predicted STK4-DDIT4L (e.g., p.Arg181Gln on STK4) and ORC4-MTUS1 (e.g., p.His166Tyr on ORC4) interfaces, respectively (Fig. 7c). Taken together, PIONEER-predicted PPI interface mutations can significantly affect drug sensitivity/resistance in antitumor treatment using both cancer cell lines and PDX models (Supplementary Table 9).

**Fig. 7.**
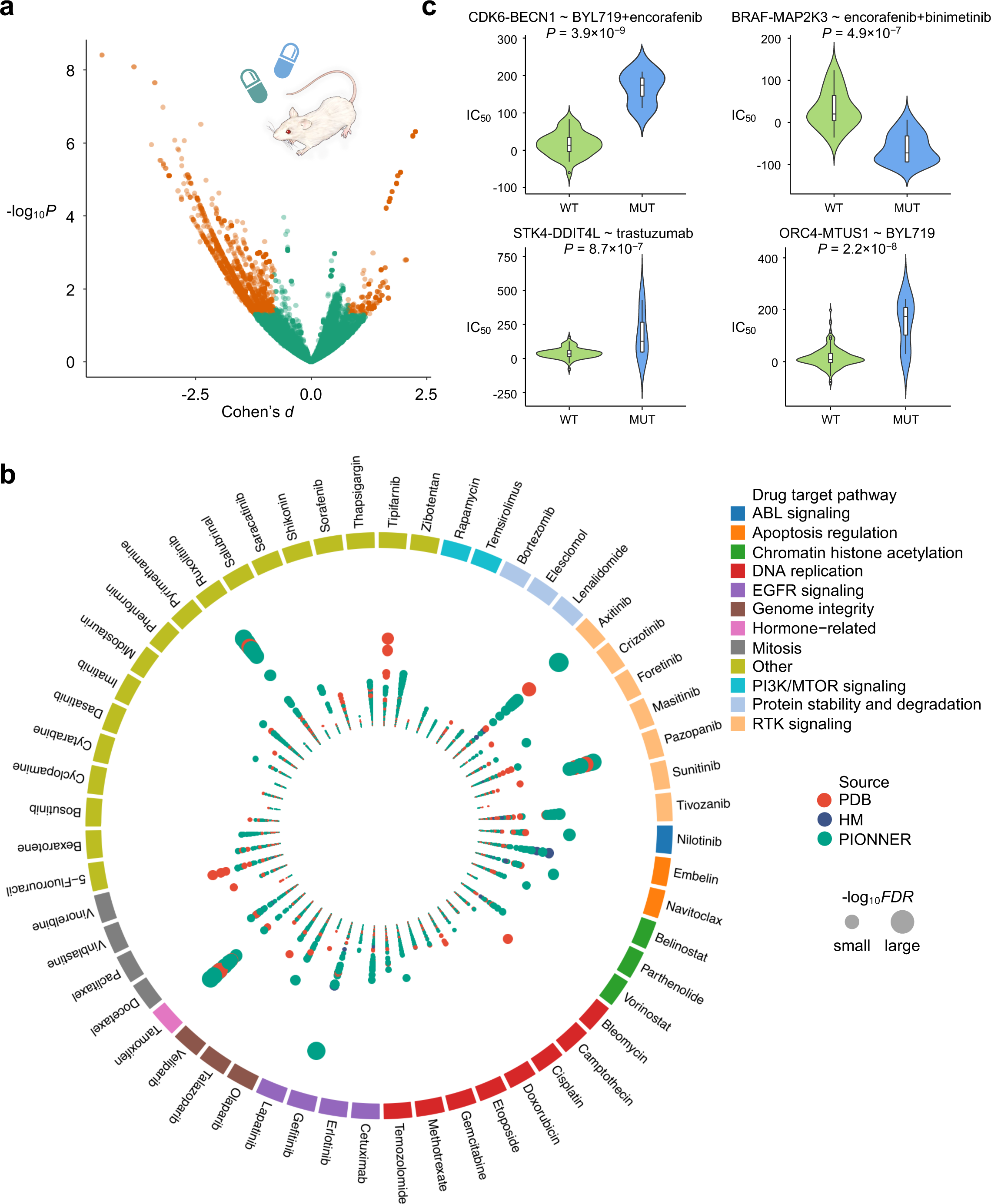
Pharmacogenomic landscape identified by the PIONEER-predicted interactome network. **a,** Drug responses evaluated by oncoPPIs in the PDX models. Effect size was quantified by Cohen’s *d* statistic using the difference between two means divided by a pooled s.d. for the data. **b,** Circos plot displaying drug responses evaluated by putative PIONEER-predicted oncoPPIs harboring a statistically significant excess number of missense mutations at PPI interfaces, following a binomial distribution across selected anticancer therapeutic agents in cancer cell lines. Each node denotes a specific oncoPPI. Node size denotes significance determined by ANOVA. Effect size was quantified by Cohen’s *d* statistic using the difference between two means divided by a pooled s.d. for the data. Node color denotes three different types of PPIs: (1) Experimental: Red; (2) HM: Blue; and (3) PIONEER: Green. **c,** Highlighted examples of drug responses. Values were calculated using ANOVA. Data are represented as a boxplot with an underlaid violin plot in which the middle line is the median, the lower and upper edges of the box are the first and third quartiles, the whiskers represent IQR × 1.5, and beyond the whiskers are outlier points. Significance was determined by ANOVA.

### Proteogenomic perturbation by PIONEER-predicted interface mutations

Recent proteogenomic study showed that somatic mutations altered protein or phosphoprotein abundance and further correlated with drug responses or cancer patient survival rate^77^. We next inspected whether PIONEER-predicted interface mutations more likely influence protein abundances in colon adenocarcinoma (COAD) and Uterine Corpus Endometrial Carcinoma (UCEC). The abundance of phosphoproteins was quantified using the tandem mass tag (TMT) assays by The National Cancer Institute’s Clinical Proteomic Tumor Analysis Consortium. We found that PIONEER-predicted interface mutations significantly reduced phosphoprotein abundances in both COAD (*P* = 0.018) and UCEC (*P* = 0.001) (Fig. 8a).

**Fig. 8.**
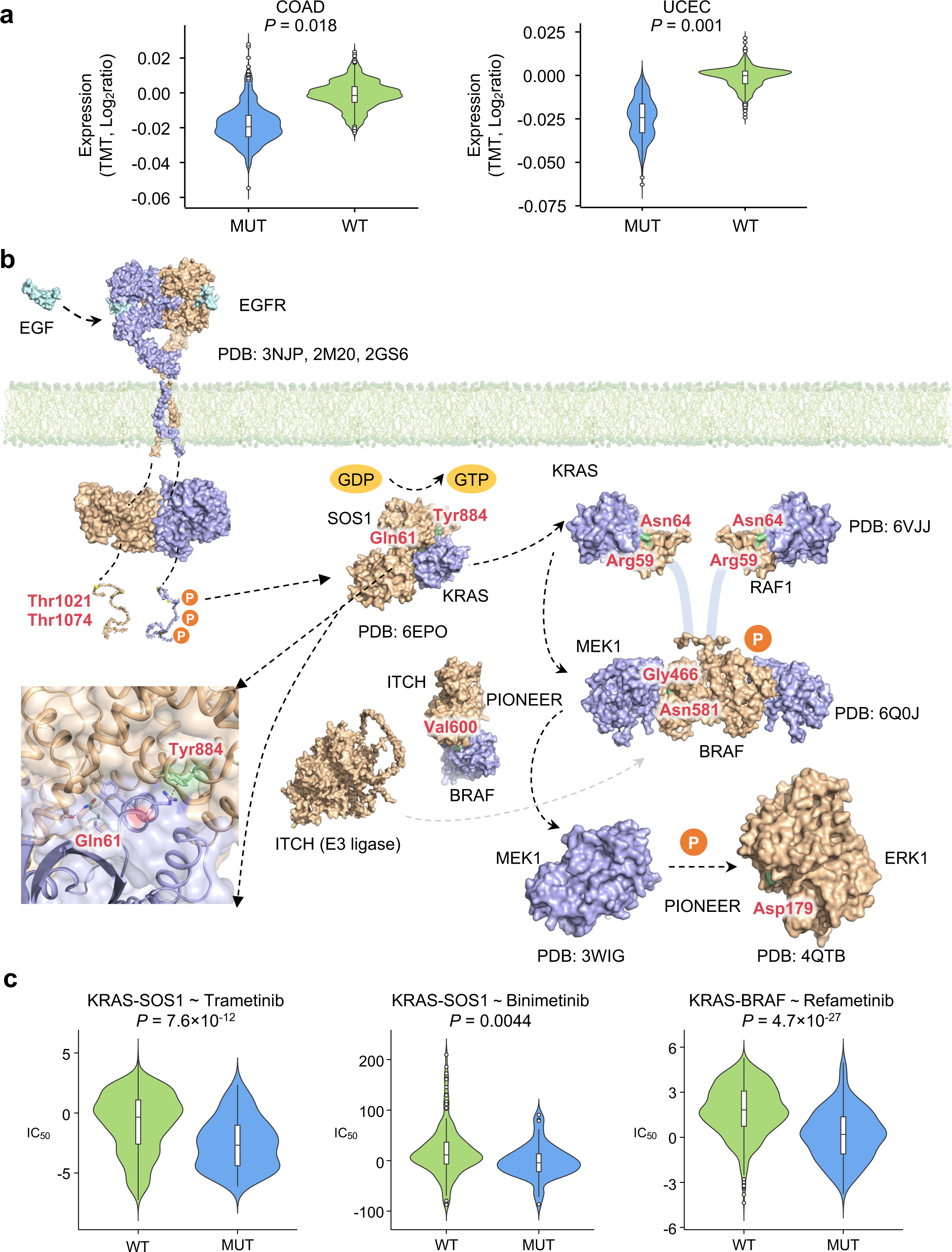
Proteogenomics of the PIONEER-predicted interactome network. **a,** Phosphorylation-associated PPI perturbing mutations altered the proteomic changes in COAD and UCEC. The abundance of proteins was quantified using the TMT technique. Significance was determined by two-tailed Wilcoxon rank-sum test. **b,** Phosphorylation-associated PPI perturbing mutations in the EGFR-RAS-RAF-MEK-ERK cascade signaling pathway. The whole transmembrane EGFR structures were constructed by three crystal structures (PDB: 3NJP, 2M20, 2GS6). The membrane model is shown in green. The phosphorylation sites are indicated by the “P” symbol. The detailed interface structure of SOS1-KRAS is also shown in the inset. The key mutated residue Gln61 on KRAS forms a hydrogen bond (purple dashed line) with residue Thr935 on SOS1, and Tyr884 on SOS1 is involved in a cation-π interaction (red dash line) with residue Arg73 on KRAS. Two subunits of RAF protein structure models were built by RAF1 and BRAF, separately (PDB: 6VJJ and 6Q0J). The two subunits are connected by a disordered loop indicated by blue cartoon lines. Two heterodimers of KRAS-RAF1 and BRAF-MEK1 constitutes the KRAS-RAF-MEK1 complex. PDB ID of each complex structure model is provided. **c,** Highlighted examples of responses to the MEK inhibitor drugs trametinib for KRAS-SOS1 mutations (n = 103 mutated cell lines, n = 827 WT cell lines), refametinib for KRAS-BRAF mutations (n = 179 mutated cell lines, n = 1,610 WT cell lines), and binimetinib for KRAS-SOS1 mutations in the PDX models (n = 62 mutated samples, n = 166 WT samples). Significance was determined by ANOVA.

We next turned to inspect how the phosphorylation-associated PPI mutations identified by PIONEER perturb EGFR-RAS-RAF-MEK-ERK signaling networks in COAD (Fig. 8b and Supplementary Table 10). The mutations involved in this signaling cascade have been suggested to regulate oncogenesis in colon and other cancers^78, 79^. EGFR dimerization was activated by EGF in the extracellular domain (PDB IDs: 3NJP and 2M20; Fig. 8b)^80, 81^. Binding of EGF triggers conformational changes in the C-terminal domain (PDB: 2GS6)^82^ and results in autophosphorylation of specific tyrosine residues, such as Tyr1068^83^. The C-terminal domain of EGFR is essential for adapter protein binding to initiate signal transduction, such as by mediating GRB2/SOS1^84^. Via PIONEER, we identified that two mutations, p.Thr1021Ile and p.Thr1074Ile on EGFR C-terminal domain, may alter the phosphorylated PPI interaction with downstream adapter protein of SOS1 (Fig. 8b). SOS1, a RAS activator by loading GTP, which was reported to allosterically interact with the Ras/Ras p.Gly12Cys mutant (PDB: 6EPO)^85^. SOS1 deficiency attenuates KRAS-induced leukemia in mouse model^86^. A selective SOS1–KRAS PPI inhibitor, BI 1701963, has been developed for advanced KRAS-mutated solid tumors in a Phase 1 clinical trial^87^. Using PIONEER, we identified two SOS1-KRAS PPI perturbation mutations, including p.Tyr884His on SOS1 and p.Gln61His on KRAS (Fig. 8b). Specifically, Tyr884 and Gln61 have strong hydrogen bond and cation-ρε interaction between KRAS and SOS1. We pinpointed that PIONEER-predicted SOS1–KRAS interface mutations are significantly related to trametinib resistance compared to WT group (*P* = 7.6×10^−12^; Fig. 8c), offering potential pharmacogenomic biomarkers for trametinib (a MEK inhibitor) in KRAS-mutant colorectal cancer^88^. Binimetinib, another MEK-selective inhibitor^89^, is significantly associated with resistance in PDX models harboring PIONEER-predicted SOS1–KRAS interface mutations (*P* = 0.0044; Fig. 8c).

GTP-bound active RAS recruits RAF proteins (e.g., RAF1 and BRAF) to the plasma membrane to orchestrate MAPK signaling^90^. Fig. 8b shows PPIs of both KRAS-RAF1 and KRAS-BRAF constructed in one structure complex. Oncogenic mutations on KRAS, such as p.Gly12Val, p.Gly13Asp, and p.Gln61Leu, are the most frequent mutations in common tumors^91^. PIONEER-predicted interface mutations of KRAS-RAF1 N-terminal, such as p.Arg59Ala and p.Asn64Ala on RAF1, are associated with a significantly reduced binding affinity of the PPI^92^, but not oncogenic mutations p.Gly12Val and p.Gly13Asp on KRAS (PDB ID: 6VJJ; Fig. 8b). In addition, we identified PIONEER-predicted KRAS-BRAF interface mutations are significantly associated with resistance of a MEK inhibitor refametinib^93^ (P= 4.7 × 10^−27^; Fig. 8c).

The key step for triggering the signaling cascade is that RAS-induced RAF dimerization subsequently phosphorylates MEK1/2 protein kinases^78^. Of RAF family members, BRAF shows the most potent activity^91^ and the BRAF p.Val600Glu mutation confers a poor survival and prognosis in colorectal cancer^94, 95^. Via PIONEER, we identified two PPI interface mutations, p.Gly466Val and p.Asn581Ser on BRAF, may mediate how BRAF coordinates MEK1 by its C-lobes in the kinase domain (Fig. 8b), consistent with a previous study^96^. Considering that the E3 ligase ITCH is also involved in BRAF regulation and binding to the kinase domain (Fig. 6a), we identified that PIONEER-predicted interface residue of Val600 on BRAF may perturb interaction between BRAF and ITCH (Fig. 8b). Phosphorylated MEK1 acts as upstream activators to phosphorylate ERK1/2 kinase activities in the MAPK cascade^97^. PIONEER-predicted interface mutation of p.Asp179Asn on ERK1 alter MEK1-ERK1 signaling network (Fig. 8b)^98^. In summary, we showed that PIONEER-predicted oncoPPIs could characterize proteogenomic alterations in the EGFR-RAS-RAF-MEK-ERK signaling pathways in colon cancer and other cancer signaling pathways if broadly applied.

### Construction of the structurally-informed interactome web server

In total, our structurally-informed interactomes cover all 282,095 experimentally-determined binary interactions in the literature for humans and seven other commonly studied organisms, including all 146,138 experimentally-determined human interactions (Figs. 1a and 2a). The web server is a user-friendly tool for genome-wide protein functional exploration through which users can identify functionally enriched areas of protein interactomes and browse multi-scale structurally-informed interactome networks (Supplementary Fig. 12). It utilizes the PIONEER framework to provide seamlessly rapid on-demand predictions for user-submitted interactions. The web tool also contains 161,244 disease-associated mutations across 10,564 disorders in HGMD and ClinVar^99^ with their per-disease enrichment pre-computed on protein interaction interfaces with 3D spatial clustering at atomic resolution for interacting protein pairs with structure models, and at the residue and domain levels for those without structures. By providing a user-friendly tool to visualize each protein and its given interactors with all available domain information, co-crystal structures, homology models, and PIONEER-predicted interfaces coupled with all known disease mutation information, PIONEER seamlessly allows users to explore the effect of mutations on 3D structures. We believe that the PIONEER web server will be instrumental in uncovering novel discoveries in relationships among these mutations with regards to the study of disease mechanisms and corresponding personalized treatment in cancer or other diseases if broadly applied.

## Discussion

The systematic characterization of structural protein-protein interactome network models from individual protein sequences and structures remains a major challenge in network medicine and is of the utmost practical and theoretical importance^2, 100^. The structural models of protein interactions can be experimentally-determined through resource-intensive and time-consuming X-ray crystallography, NMR, and more recently, cryo-EM experiments. Computational methods, such as docking and homology modeling, have been developed to predict PPI interfaces. Although docking methods can achieve atomic resolution, they do not generally behave well especially in cases where proteins undergo large conformational changes^101, 102^. Homology modeling, which covers only another ∼3% of interactions, infers protein interfaces from templates of homologous complexes^13^. To date, ∼91% of PPIs still do not have structural information. Hence, most network studies model proteins as graph-theoretical nodes, which ignores the structural details of the individual proteins as well as the spatial constraints of their interactions.

We previously developed the innovative ensemble random forest-based pipeline, ECLAIR, which was among the first to generate the full-proteome structural human interactome network. Over the last several years, deep learning techniques have shown exceptional representation learning abilities and much improved performance, especially in processing 3D protein structures^103^, compared to traditional machine learning algorithms, including random forest. A number of methods^21, 25, 26^, therefore, have been proposed that have gained considerable traction for protein interface prediction. However, a major limitation of these methods is that they do not leverage the notable combination of feature representations indicated in both structures and sequences well nor do they effectively integrate embeddings from both the local regions of target residues and the whole proteins, thereby leaving much room for improvement in prediction quality. Recently, RoseTTAFold^104^ and AlphaFold2^105^ have made a significant breakthrough in generating single protein structural models from solely protein sequences, which is extremely helpful for protein interactome studies. Even more recently, AlphaFold-based methods, such as AlphaFold-Multimer and AF2Complex, have been developed to generated structural models for multi-chain protein complexes. AlphaFold Multimer requires joint multiple sequence alignments paired by species while AF2Complex does not, but they are both very computationally intensive, and, thus, cannot be scaled to generate models for the hundreds of thousands of interactions in whole interactomes. Furthermore, a recent study has shown that AlphaFold2 can successfully generate high-quality models for only ∼2% of human interactions without known homologous structures^106^.

Here, we present a comprehensive, multiscale structurally-informed interactome framework and web server, which we named PIONEER, to combine seamlessly genomic-scale data with structural proteomic analyses. This resource is based on our deep-learning-based ensemble framework, which accurately predicts partner-specific interaction interfaces for all protein-protein interactions in humans and seven other common model organisms. We found PIONEER outperforms other existing state-of-the-art methods, including our previously developed method, ECLAIR. Moreover, large-scale statistical analysis and mutagenesis experiments show that PIONEER-predicted interfaces reveal similar biological significance as those of known interfaces in the PDB. Further analysis shows that interactome PIONEER plays a pivotal role in dissecting the pathobiology of diseases. Our work is implemented as both a web server platform and a software package to help facilitate systematic structural analysis in genomic studies, thereby allowing the wider scientific community to adopt and further develop upon our PIONEER framework.

With the rapid advances in sequencing technologies, multiple whole genome/exome sequencing projects are currently being carried out, including TCGA, cardiovascular medicine (i.e., National Heart, Lung and Blood Institute’s Trans-Omics for Precision Medicine program^107^), and Alzheimer’s disease sequencing project^108^, to identify trait/disease-associated genes and mutations. Although the 3D structure of each protein fundamentally determines its function, systematic structural analysis is currently not part of any major genomic study pipeline. One of the main reasons for this disconnection is the poor coverage of atomic-resolution structural models of proteins and their interactions. We expect our comprehensive structurally-informed interactomes generated by PIONEER, which provides high-quality partner-specific interfaces on the scale of the whole interactome, will help bridge the gap between genome-scale data and structural proteomic analyses. With the high-quality and comprehensive map of protein interfaces, there are numerous valuable extensions considering the biophysical effects induced by mutations at protein interfaces, such as the investigation of disease etiology and the corresponding drug prioritization, and prediction of specific disease pathobiology. The partner-specific property of PIONEER-generated structurally-informed interactomes also allows us to study the pleiotropic effects of genes. Therefore, the powerful and comprehensive PIONEER framework will make such extensive research possible, and, more importantly, provide potentially unforeseen avenues for drug design and therapeutics.

The current experimentally-determined binary human interactome in the literature is far from complete. The scientific community has dedicated extensive efforts both experimentally (such as HuRI^109^, BioPlex^110^, and OpenCell^111^) and computationally (such as PrePPI^112^ and HIGH-PPI^113^), to ascertaining which pairs of human proteins interact. As more protein interactions are detected for human proteins, our PIONEER resource will be regularly updated to make interface predictions for newly-released human PPIs. Specifically, PrePPI is a very interesting method that uses structural homology to make accurate PPI predictions. Although PrePPI does not make interface predictions (thus not comparable to PIONEER), the structural information obtained by PrePPI can be incorporated into our PIONEER pipeline as an additional feature to potentially improve our interface prediction performance in the future.

We have implemented PIONEER and the resulting multiscale structurally-informed interactomes into a user-friendly web server platform, and also constructed the PIONEER framework as a flexible software package, which benefits users and developers as well. We demonstrated PIONEER’s utility in discovering new biological insights into multiple genome medicine studies. We combined PIONEER predictions along with co-crystal structures and homology models to reconstruct a subnetwork in the human interactome that is enriched with disease mutations. Specifically, we demonstrated the PIONEER-predicted interface mutations significantly enriched in both somatic cancer and germline mutations. Moreover, PIONEER-predicted interface mutations highly correlated with cancer patient survival rate and anti-cancer responses in both tumor cell lines and PDX models. In addition, we experimentally validated PIONEER-predicted PPI interface mutations using CCND1-CDK4 and KEAP1-NRF2 as two examples. In particular, PIONEER-predicted PPI interface mutation altering E3 ligase between significantly promoted non-small cell lung cancer cell growth. In summary, these comprehensive observations illustrated high clinical utility of the PIONEER-predicted structurally-informed interactome networks in genome research and precision medicine studies.

The web server platform allows for a broad range of investigations related to protein interfaces on a genome-wide scale while also carrying out on-demand interface predictions for user-uploaded interactions. Furthermore, the software package increases the utilization, maintenance, and further development of PIONEER by the wider scientific community. The accompanying web portal, containing all relevant PDB structures and homology models, as well as predicted interface residues and domains, significantly reduces the barrier-to-entry to perform systematic structural analysis for most genetics/genomics researchers, and allows for implementing such analyses (e.g., looking for enrichment of mutations on protein structures and interfaces at various resolutions, distinguishing mutations in different interfaces of the same protein) in genome/exome sequencing projects. Additionally, the continually updated and extendable software package bears great potential for stimulating the adoption and development of our PIONEER framework for related genome medicine research.

## Methods

### PPI interface data construction

We compiled 282,095 binary interactions for *H. sapiens, A. thaliana, S. cerevisiae, D. melanogaster, C. elegans, M. musculus, S. pombe* and *E. coli* in total, including 9,123 full experimentally-determined binary interactions in humans. The interactions with known co-crystal structures in the PDB were used to form the training, validation, and testing datasets to build PIONEER models, which then predicted the interfaces for all the other interactions without known co-crystal or homologous structures. The homologous structures are collected from Interactome3D^114^.

We calculated the partner-specific interface residues for those interactions with known co-crystal structures in the PDB. SIFTS^115^ was then used to map the UniProt-indexed residues to the PDB-indexed residues. To determine the interface residues, we used NACCESS^116^ to assess the change in solvent-accessible surface area of the protein in complex and in isolation. Specifically, an interface residue is defined as a residue that is a surface residue (≥15% exposed surface) with its SASA decreasing by ≥1.0Å^2^ in the complex. We reviewed all available structures in the PDB for an interaction, and considered a residue to be in the interface of that interaction if it had been calculated to be an interface residue in at least one of the corresponding co-crystal structures. By only considering interactions for which aggregated co-crystal structures have been combined to cover at least 30% of the UniProt residues for both interacting proteins, the training, validation, and independent benchmark testing datasets were built, which included a random selection of 2,615, 400, and 400 interactions with known co-crystal structures, which include 1,191,036; 174,739; and 186,326 residues for sufficient model training, validation, and testing, respectively. The number of positive residues (interface residues) compared to negative residues (non-interface residues) in this dataset is 175,911/1,015,125 (17.3%), 25,641/149,098 (17.2%), and 27,744/158,582 (17.5%), respectively. It is important to note that a single residue may be labelled as positive for a specific interaction, and labelled negative for other interactions. Additionally, we ensured that no homologous interactions or repeated proteins existed between any of the two datasets to guarantee the robustness and generalizability of our models, and a fair performance evaluation. We define homologous interactions as a pair of interactions where both proteins in one interaction are homologs of both proteins in the other interaction. Three iterations of PSI-BLAST^117^ at an E-value cutoff 0.001 were carried out to determine the protein homologs.

### Feature characterization

Our previous pipeline ECLAIR employed a set of representative feature groups to describe the residues, including biophysical residue properties, evolutionary sequence conservation, co-evolution, SASA, and docking-based metrics. While retaining all features from ECLAIR here, we also implemented two new feature groups to seek a more comprehensive and in-depth feature characterization. From each feature group, we synthesized a variation of features using scaling, by which we mean that each feature used its raw calculated values and normalized values against the average of all positions per protein.

#### 1. Predicted protein structural properties

Here, in addition to SASA used in ECLAIR, we also used RaptorX^118^ to predict the solvent accessibility state (buried, medium, and exposed) and the secondary structure state (helix, strand, and coil), which are especially useful for proteins without structural information. The predicted solvent accessibility and secondary structure were represented by their predicted probabilities.

#### 2. Pair potential

In our previous ECLAIR, we used co-evolution and docking-based features to describe this partner-specific issue, which, however, is not available for the interactions not having co-evolution and structural information, respectively. Although our deep learning architectures solve this problem by considering the partner protein embedding derived from the inputs, here, inspired by the pairing preferences at protein-protein interfaces proposed by Glaser et al.^119^, we also proposed a novel feature, pair potential, to specifically characterize the interface propensity for the target protein with respect to its respective partner. Pair potential was developed solely based on sequence information so it can be applied to all the binary protein interactions. To calculate pair potential, we need first to calculate the propensity matrix *G* of contacting residue pairs:

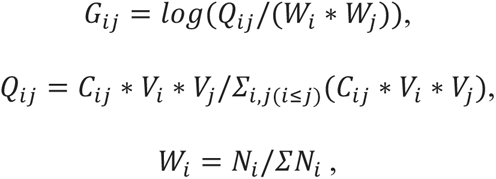

where *Q_ij_* is the normalized number of contacting residue pairs between residues *i* in the target protein and *j* in the partner protein, *C_ij_* is the number of contacting residue pairs observed between residues *i* and *j*, *V_i_* is the volume of residue *i*, *W_j_* is the normalized frequency of interfaces, and *N_i_* is the number of interface residues *i* from both target and partner proteins. The propensity matrix *G* was calculated using only interface residues in our training dataset. The pair potential for each residue in the target protein can, thus, be calculated as:

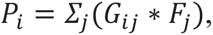

where *F_j_* is the frequency of residue *j* in the partner protein.

#### Model building

In order to ensure that every residue is meticulously predicted through the maximal amount of available information from both proteins in an interaction pair, we built four deep learning models in which each model takes different interactions as input based on the availability of structures.

1. Structure-Structure model (Fig. 1c and Supplementary Figs. 1a and 2): For interactions where both proteins have structural information available, the structure and sequence information were embedded through GCNs with ARMA filters and bidirectional RNNs with GRUs, respectively. Specifically, GCN utilizes the structural information from graph representations of protein structures where each node represents a residue, and each edge signifies that two residues are adjacent. For each node, GCN incorporates its spatial neighborhood information to generate a more comprehensive residue representation whereas RNN explores amino acid sequences to include the sequential neighborhood information of each residue. The RNN extracts the upstream and downstream sequence information from each residue. Through the concatenation and mean aggregation, the residue embeddings of both target protein and partner protein were then converted to protein embeddings, respectively. Finally, the residue embeddings, target protein embedding, and partner protein embedding were concatenated and fed into the fully connected layers to make prediction for each residue in the target protein.
2. Sequence-Sequence model (Fig. 1d and Supplementary Figs. 1b and 2b): For interactions where neither protein has structural information, the sequence information of both proteins was fed into the RNNs. Next, in a manner similar to that described in the Structure-Structure model, the residue embeddings, target protein embedding, and partner protein embedding were concatenated and fed into the fully connected layers to make prediction for each residue in the target protein.
3. Structure-Sequence and Sequence-Structure models (Fig. 1e,f and Supplementary Figs. 1c,d and 2): The use of Structure-Sequence or Sequence-Structure model depends on whether target protein or partner protein has structural information, respectively. The transfer learning was used in these two models, which means the pre-trained GCNs-RNNs in the above Structure-Structure model and RNNs in the above Sequence-Sequence model were deployed in the Structure-Sequence model and the Sequence-Structure model for the processing of proteins with and without structural information, respectively. Subsequently, in a manner similar to that described in the Structure-Structure model and Sequence-Sequence model, the residue embeddings, target protein embedding, and partner protein embedding were concatenated, and fed into the fully connected layers to make prediction for each residue in the target protein.

We compiled a set of representative protein structures from the PDB and ModBase^120^ for each protein. For ModBase models, we only consider the models with a ModPipe Quality Score (MPSQ) ≥1.1. The structures were sorted by the coverage of UniProt residues based on SIFTS, excluding any homologous PDB structures of interacting protein pairs. Each residue in a target protein was then reviewed if it has structural information; if so, it was predicted using that protein’s first corresponding structure which contains the structural information of that residue; otherwise, it was predicted using the sequence information. For the partner protein which has structural information, we only used the corresponding structure with the highest UniProt coverage. To make our tool more practically useful and to avoid the memorization of known interfaces, we use the single protein structure which is not from co-crystal or homologous co-complex structures to train the model.

Our PIONEER framework was implemented using PyTorch. To maximize model performance, we carried out comprehensive hyperparameter optimization for the neural network architectures, and the optimal set of hyperparameters were determined by maximizing area under the receiver operating characteristic (AUROC) on the validation set. All four models were trained with cross entropy loss and the Adam optimizer; the kernel activation function^121^ was used in GCNs and fully connected layers. The hyperparameters used for these four models can be found in our accompanying PIONEER software package. To solve the variable length inputs, we trained all four models in a mini-batch mode with only a single protein pair.

### Performance evaluation

After identifying the best hyperparameters for each model, a thorough examination was performed using the benchmark testing set. Models were ordered based on their AUROCs on the validation set, which means the priorities of models are Structure-Structure, Structure-Sequence, Sequence-Structure, and Sequence-Sequence, respectively. For the overall performance, the raw prediction score of each residue was taken from the results of the model with highest priority according to the availability of structures of the target protein containing that residue and its partner protein. We further compared PIONEER with a number of existing state-of-the-art methods, including ECLAIR, PeSTo, ScanNet, BIPSPI+, MaSIF-site, DeepPPISP, SASNet, PIPGCN, DELPHI^122^, SCRIBER^123^, and DLPred^124^. We also reported performance metrics at various discrete and comparable levels of confidence, which consist of Very low, Low, Medium, High, and Very high prediction categories, by evenly separating into fifths our raw prediction scores.

### Interface prediction

By further incorporating AlphaFold2-predicted structures, we predicted interface residues for the remaining 256,946 interactions not resolved by either PDB structures or homology models. Each residue was then predicted by the model of the ensemble with the highest priority according to the availability of structures of the target protein containing that residue and its partner protein.

### Mutagenesis validation experiments

We performed mutagenesis experiments where we introduced random human population variants from the Exome Sequencing Project^125^ into predicted interfaces, known interfaces, and non-interfaces. We randomly selected mutations of predicted interfaces in each of the PIONEER prediction categories (very low-very high). We also selected random mutations of known interfaces and non-interfaces in co-crystal structures in the PDB as positive and negative controls. The selected mutations were introduced into the proteins according to our Clone-seq pipeline^37^. We generated 2,395 mutations on 1,141 proteins and examined their impact on 6,754 mutation-interaction pairs (either disrupting or maintaining the interactions) using our high-throughput Y2H assay.

### Y2H assay

Y2H was performed as previously described^6^. Gateway LR reactions were used to transfer all wild-type/mutant clones into our Y2H pDEST-AD and pDEST-DB vectors. All DB-X and AD-Y plasmids were transformed into the Y2H strains MATα Y8930 and MATa Y8800, respectively. Thereafter, each of the DB-X MATα transformants (wild-type and mutants) were mated with corresponding AD-Y MATa transformants (wild-type and mutants) individually through automated 96-well procedures, including inoculation of AD-Y and DB-X yeast cultures, mating on YEPD media (incubated overnight at 30 °C), and replica-plating onto selective Synthetic Complete media lacking histidine, leucine, and tryptophan, and supplemented with 1 mM of 3-amino-1,2,4-triazole (SC-Leu-Trp-His+3AT), SC-Leu-His+3AT plates containing 1 mg/l cycloheximide (SC-Leu-His+3AT+CHX), SC-Leu-Trp-Adenine (Ade) plates, and SC-Leu-Ade+CHX plates to test for CHX-sensitive expression of the LYS2::GAL1-HIS3 and GAL2-ADE2 reporter genes. The plates containing cycloheximide were used to select for cells that do not have the AD plasmid due to plasmid shuffling. Spontaneous auto-activators^126^, therefore, were identified by growth on these control plates. These plates were incubated overnight at 30 °C and “replica-cleaned” the following day. Subsequently, plates were incubated for three more days, after which positive colonies were scored as those that grow on SC-Leu-Trp-His+3AT and/or on SC-Leu-Trp-Ade, but not on SC-Leu-His+3AT+CHX or on SC-Leu-Ade+CHX. Disruption of an interaction by a mutation was defined as at least 50% reduction of growth consistently across both reporter genes when compared to Y2H phenotypes of the corresponding wild-type allele as benchmarked by 2-fold serial dilution experiments. All Y2H experiments were repeated 3 times.

### Co-immunoprecipitation

The first co-immunoprecipitation assay was conducted to validate the PIONEER partner-specific interface prediction. Specially, HEK293T cells were maintained in DMEM medium supplemented 10% Fetal Bovine Serum. Cells were seeded onto 10 cm dishes and incubated until 40-50% confluency, and were transfected with a mixed solution of 3 μg of bait construct (CCND1), 3 μg of prey construct (CDK4 or TSC2), 30 μl of 1 mg/ml PEI (Polysciences, 23966), and 1.2 ml OptiMEM (Gibco, 31085-062). After 48 hrs incubation, transfected cells were washed three times in 10 ml DPBS (VWR, 14190144), resuspended in 500 μl NP-40 lysis buffer (50 mM Tris pH 7.5, 150 mM NaCl, 5 mM EDTA, 1.0% NP-40) and incubated on the ice for 30 min. Whole lysate is sonicated on a sonifier cell disruptor (BRANSON, 500-220-180) for 120 sec at 40% amplitude. Extracts were cleared by centrifugation for 15 min at 16,100 g at 4 °C. For co-immunoprecipitation, 500 μl cell lysate per sample reaction was incubated with 15 μl of EZ view Red Anti-FLAG M2 Affinity Gel (Sigma, F2426) overnight 4 °C with a nutator. After incubation, bound proteins were washed three times in NP-40 lysis buffer and then eluted in 200 μl of elution buffer (10 mM Tris-Cl pH 8.0, 1% SDS) at 65 °C for 15 min. FLAG-co-purified samples were run on 8% SDS-PAGE gel and the proteins were transferred to PVDF membranes. Anti-FLAG (Sigma, F1804), and Anti-MYC (Invitrogen, 132500) at both 1:5,000 dilutions were used for immunoblotting analysis.

We also validated mutation effects for KEAP1-NRF2 by co-immunoprecipitation assay, in which HEK293T cells were co-transfected with WT KEP1 expressing vector and either WT NRF2, p.Thr80Lys, p.Glu79Lys, or p.Leu30Phe expressing vectors for 48 hours. Cells were lysed with NP-40 lysis buffer on ice, and supernatants were incubated with anti-HA antibody coupled with protein A/G beads (Santa Cruz,) overnight. Immunoprecipitated complexes were washed with NP-40 lysis buffer for 3 times, and were then eluted and subjected to Western blotting.

### Collection and preparation of genome sequencing data

We collected variant data across multiple sources including TCGA, MSK-MET, 1KGP, ExAC, HGMD, Cancer Cell Line Encyclopedia and genomic profiling of PDXs from previous study^76^. For unannotated datasets, we used VEP^127^ to annotate these variants in order to identify the corresponding amino acid changes. We regard one PPI as mutated if one variant affects the amino acid residue in the interface of either protein involved in the interaction.

### Significance determination of PPI interface mutations

The significance of PPI interface mutations were tested using the method as described in our previous study^8^. A PPI in which there is significant enrichment in interface mutations in one or the other of the two protein-binding partners across individuals will be defined as an oncoPPI. For each gene *g_i_* and its PPI interfaces, we assume that the observed number of mutations for a given interface follows a binomial distribution, binomial (*T*, *p_gi_*), in which *T* is the total number of mutations observed in one gene, and *p_gi_* is the estimated mutation rate for the region of interest under the null hypothesis that the region was not recurrently mutated. Using length(*g_i_*) to represent the length of the protein product of gene *g_i_*, for each interface, we computed the *P* value - the probability of observing >k mutations around this interface out of *T* total mutations observed in this gene - using the following equations:

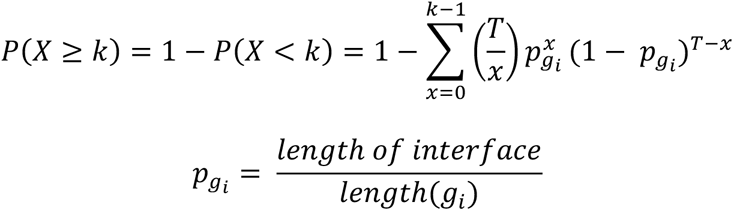

Finally, we set the minimal *P* value across all the interfaces in a specific protein as the representative *P* value of its coding gene *g_i_*, denoted *P*(*g_i_*). The significance of each PPI is defined as the product of *P* values of the two proteins (gene products). All *P* values were adjusted for multiple testing using the Bonferroni correction.

### PPI system construction of E3 ligases

355 E3 ubiquitin ligases were retrieved and merged from E3net and Ubinet2.0. 4,613 E3s-associated oncoPPIs were analyzed after removing PPIs with homodimers or without gene name. These oncoPPIs include 198 oncoPPIs from the PDB database, 197 from Homology Models, and 4,218 from PIONEER. The correlations between mutations in E3 oncoPPIs and anticancer drug responses in TCGA cell lines, PDX models, and cancer survival rates from TCGA and MSK MetTropism datasets were then calculated (Supplementary Table 9).

Complex crystal structures (PDBs: 3HQH, 3O37, 4O1V, 5G05, 5LNB, 5VZU, 6F8F, 6I68, 6M90, 6WCQ, 7K2M and 7LIO) were accessed from the RCSB PDB protein data bank. The structures in the complex without co-crystal structures were retrieved from the AlphaFold2 portal (https://alphafold.ebi.ac.uk). PIONEER predicted PPI models were constructed using HADDOCK^128^. The names, mutations and PDB ID were also shown in Supplementary Table 8.

### The linear ANOVA model

We used the drug response data of human cancer cell line from GDSC datasets and to investigate the association of PPI interface mutation with drug response. For each drug, a drug-response vector consisting of IC_50_ values was modeled using the status of a genomic feature (whether a PPI interface is mutated), the tissue of origin of the cell lines, screening medium, and growth properties by fitting a linear model. A genomic feature-drug pair was tested only if the final IC_50_ vector contained at least 10 positive cell lines. The effect size was quantified through Cohen’s *d* statistic using the difference between two means divided by a pooled standard deviation for the data. The resulting *P*-values were corrected by the Benjamini-Hochberg method^129^. Similar to cell line drug response analysis, we also used the drug response data from high-throughput screening using PDX models to study the association of PPI interface mutation with drug response using linear model. All statistical analyses were performed using the R package (v4.2.0, http://www.r-project.org/).

### System structural construction of EGFR-RAS-RAF-MEK-ERK signaling network

EGR-EGFR complex were constructed by three crystal structures (PDB: 3NJP, 2M20, 2GS6). Membrane models were built by CHARMM-GUI^130^. SOS1-KRAS complex (PDB: 6EPO), KRAS-RAF1 complex (PDB: 6VJJ), MEK1-BRAF complex (PDB: 6Q0J), MEK1 (PDB: 3WIG), and ERK1 (PDB: 4QTB) were accessed from the RCSB PDB protein data bank. Two subunits of RAF proteins are represented by RAF1 and BRAF, separately. The ITCH-BRAF complex model was generated using HADDOCK. All images were processed using PyMOL. The complex names, mutations and PDB ID were also shown in Supplementary Table 10.

### Cell viability assay

H1975 cells were transfected with NRF2 T80K/WT-expressing vectors or empty vectors using Lipofectamine 3000 (Thermofisher). For growth curve measurement, 3,000 cells were planted into 96-well plates, and viability was measured using CellTiter 96 AQueous MTS Reagent (Promega,) at days 0, 2 and 4.

### Colony formation assay

For the colony formation assay, 2,000 cells were plated into 6-well plates. After 2 weeks, cells were fixed with 4% paraformaldehyde and stained with crystal violet. Relative growth index was analyzed using ImageJ software^131^.

### Web server development

The PIONEER web server is written in Python using the Django framework. The web server displays the full human interactome using co-crystal structures from PDB, homology models from Interactome3D^114^, and PIONEER-predicted interfaces. These data, along with auxiliary data about each protein, are stored on a Postgres database. The front-end displays this data using jQuery in two different web pages. The first page displays all information associated with a single protein-protein interaction. We display the protein-protein interaction interface information, show where the interface residues lie on the PFAM domains for each protein, and depict where the disease mutations are enriched on the interaction interface. The disease mutation information is obtained from HGMD. We display a number of 3D structures to help visualize the interaction interface. These includes the high-quality 3D structures from PDB and ModBase associated with each protein as well as any available high-quality PDB co-crystal structures. Furthermore, we display all AlphaFold structures associated with each protein. These structures are displayed using the MichelaNGLo viewer^132^. The second webpage displays a subset of the human interactome associated with a particular protein or a particular disease, depending on the user’s input on the homepage. We use the springy.js network visualization library here.

The web server also hosts interaction data for eight species: *H. sapiens, A. thaliana, S. cerevisiae, D. melanogaster, C. elegans, M. musculus, S. pombe* and *E. coli*. Most importantly, the PIONEER web server also consists of an on-demand prediction pipeline that allows users to predict the interaction interface between any two proteins given their UniProt IDs. The on-demand prediction pipeline returns the raw predictions made by PIONEER and shows the two-dimensional protein structure visualizing which regions of the protein are more likely to lie on the interaction interface. The prediction pipeline utilizes the SLURM job scheduler to serve these requests.

### Data availability

Mutation data from the TCGA study were downloaded from NCI genomic data commons (https://portal.gdc.cancer.gov). MSK MetTropism dataset was downloaded from cBioPortal (https://www.cbioportal.org/study/summary?id=msk_met_2021). Variant data from 1000 Genomes Project were downloaded from NCBI FTP Site (ftp://ftp-trace.ncbi.nih.gov/1000genomes/ftp). The ExAC data set was downloaded from gnomAD (https://gnomad.broadinstitute.org/downloads#exac-variants). Variants collected by HGMD were downloaded from (https://www.hgmd.cf.ac.uk/ac/index.php). Genomic variants and drug response data of human cancer cell lines were downloaded from GDSC datasets (https://www.cancerrxgene.org/downloads/bulk_download). Genomic profiling of PDXs and drug response curve metrics of PCTs were downloaded from the Supplementary Table 1 of the corresponding paper (https://www.nature.com/articles/nm.3954#Sec28). All other data supporting the results in this study are available in supplementary materials, and at https://pioneer.yulab.org.

### Code availability

The source code of PIONEER is available at https://github.com/hyulab/PIONEER.

## Acknowledgments

This work was supported by the National Institute of General Medical Sciences (R01GM124559, R01GM125639, R01GM130885, and RM1GM139738) and The National Institute of Diabetes and Digestive and Kidney Diseases (R01DK115398) to H.Y.; the National Institute on Aging under Award Number R01AG084250, R56AG074001, U01AG073323, R01AG066707, R01AG076448, R01AG082118, RF1AG082211, and R21AG083003, and the National Institute of Neurological Disorders and Stroke (NINDS) under Award Number RF1NS133812 to F.C.; and the National Human Genome Research Institute (U01HG007691), the National Heart, Lung, and Blood Institute (R01HL155107, R01HL155096, and U54HL119145), the American Heart Association (AHA957729), and European Union Horizon Health 2021 (101057619) to J.L. C.E. is the Sondra J. and Stephen R. Hardis Chair of Cancer Genomic Medicine at the Cleveland Clinic.

## Author contributions

D.X., Y.Q., J.Z., Y.Z., D.L., F.C. and H.Y. conceived and developed the project. D.X., D.L. and S.L. developed the models and conducted computational experiments. S.G., M.T. and S.L. built the web server. W.L., J.K. conducted biological experiments. D.X., Y.Q., J.Z., Y.Z., D.L., F.C. and H.Y. performed the analyses. D.X., Y.Q., J.Z., Y.Z., D.L., S.G., C.E., J.L., F.C. and H.Y. wrote and critically revised the manuscript. All authors discussed the results and reviewed the manuscript.

## Supplementary Data

**Data 1.** The labeled dataset used for PIONEER model training, validation, and testing.

**Data 2.** List of somatic mutations in 33 cancer types of TCGA that affect PPI interfaces.

**Data 3.** Significance test of somatic mutation enrichment in PPI interfaces by 33 cancer types in TCGA.

